# Cascaded Processing of Amplitude Modulation for Natural Sound Recognition

**DOI:** 10.1101/308999

**Authors:** Takuya Koumura, Hiroki Terashima, Shigeto Furukawa

## Abstract

Temporal variation of sound envelope, or amplitude modulation (AM), is essential for auditory perception of natural sounds. Neural representation of stimulus AM is successively transformed while processed by a cascade of brain regions in the auditory system. Here we sought the functional significance of such cascaded transformation of AM representation. We modelled the function of the auditory system with a deep neural network (DNN) optimized for natural sound recognition. Neurophysiological analysis of the DNN revealed that AM representation similar to the auditory system emerged during the optimization. The better-recognizing DNNs exhibited larger similarity to the auditory system. The control experiments suggest that the cascading architecture, the data structure, and the optimization objective may be essential factors for the lower, middle and higher regions, respectively. The results were consistently observed across independent datasets. These results suggest the emergence of AM representation in the auditory system during optimization for natural sound recognition.

## Introduction

Natural sounds such as speech and environmental sound exhibit rich patterns of amplitude envelope (Fig. 1a). Temporal variation of amplitude envelope, called amplitude modulation (AM), is one of the most important physical dimensions for auditory perception^1,2^. Humans can recognize speech contents and identify daily sound based on its AM patterns even if its fine temporal structure is substantially deteriorated^3,4^. AM patterns of a sound is usually characterized by their frequency components, AM frequencies (Fig. 1b).

**Fig. 1.**
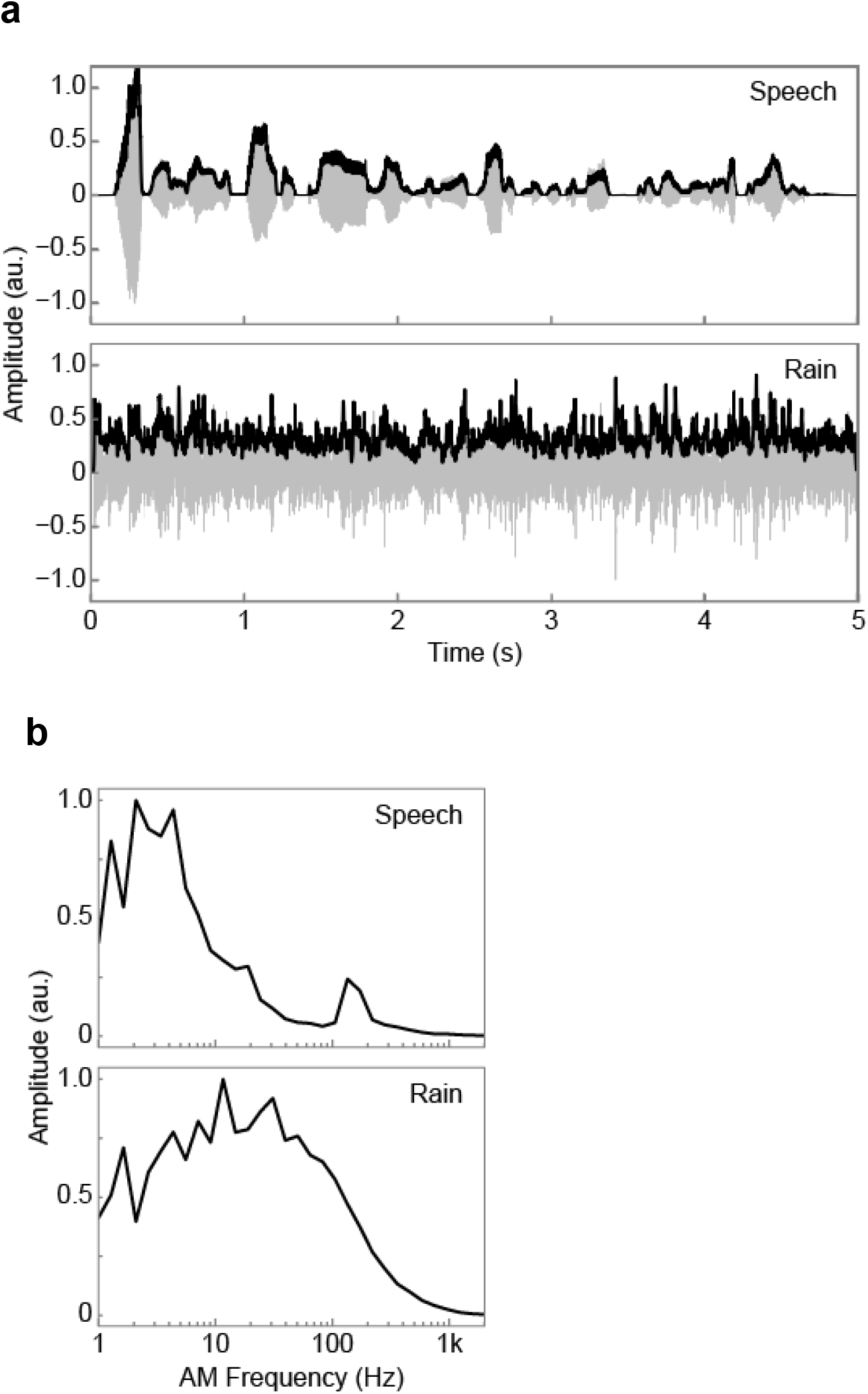
Rich repertoires of amplitude envelope in natural sounds. (a) Examples of sound waveforms (grey) and their amplitude envelopes (black) of natural sound. Sounds of speech (top) and rain (bottom) are shown. Amplitude envelopes of speech and rain appeared different. (b) Modulation spectra, distributions of the AM frequency components, of the sounds in (a). The modulation spectrum was calculated as the root mean square of the filtered envelope with a logarithmically spaced bandpass filter bank. Each modulation spectrum is normalized by its maximum. The lower and the upper peak in the modulation spectrum of speech (top) probably contain the information of the speech content and the speaker, respectively. The modulation spectrum of the rain sound (bottom) appeared different from the one of the speech.

Perceptual importance of AM has driven physiologists to seek neural representation of AM in the auditory system. The auditory system converts physical properties of a sound stimulus to neural activities and transmit them through a cascade of brain regions for further processes of perception^5,6^. In the auditory system, not only do some neurons fire synchronously to the stimulus amplitude envelope, tuning to the AM frequency is also observed both in the degree of spike synchronization and in the spike rate. This implies that the auditory system performs some kinds of frequency analysis in the AM domain by temporal and rate coding^7^, which means AM coding with spike temporal patterns and average spike rate, respectively. A range of studies broadly agree that the characteristics of AM coding change somewhat systematically along the processing stages from the periphery to the cortex^5,7^. Along the auditory neuraxis, the AM frequency to which neurons synchronize gradually decreases, and the number of neurons which performs rate coding of AM frequency gradually increases. The latter phenomenon is called temporal-to-rate conversion.

An ever-growing number of physiological studies are conducted for various brain regions and animal species to expand the dataset. There are also experimental and theoretical studies that attempt to explore neural mechanisms that may realize the above observations^8–12^. Those approaches have revealed how the system works. However, they do not answer why the system has to be organised in that way. We would like to ask the functional significance of the systematic transformation of AM representation through the cascade of regions. Is it a consequence of evolution for efficiently extracting essential signals from natural sounds for survival, or merely a byproduct of other biological constraints?

### A deep neural network as a functional model of sensory systems

For explaining functional significance in several sensory modalities or dimensions^13–16^, machine learning techniques have been proven to be effective. Modelling with such techniques are generally not heavily based on anatomical or physiological assumptions, but the architectures and the parameters of the model can be trained to process natural stimuli for ethologically relevant objectives. Thus, the trained model is expected to express effective representation of natural stimuli for such objectives, and if the representation is similar to that observed in an actual sensory system, it is highly likely that the sensory system is also adapted to effectively processing sensory information for survival.

A deep neural network (DNN) is one of the most successful machine learning techniques both for automatic data processing^17–19^ and for explaining neural representation of sensory information^20–24^. A DNN consists of multiple layers with multiple units, and a unit in a layer integrates outputs of the units in the lower layer and sends outputs to other units in the upper layer. Apart from this, the DNNs in the previous sensory studies are neither designed to reproduce any physiological or anatomical properties of the biological neurons, nor optimized to specific neural activities. Nevertheless, DNNs trained on natural recognition tasks outperform other conventional carefully-designed models in predicting the neural activities.

In this study, we trained a DNN to estimate categories of non-human natural sound consisting of animal vocalizations and environmental sounds. The task is to classify 0.19-s long sound waveforms into one of 18 categories. Our DNN takes raw data (that is, amplitude waveform) as an input and estimates the category of the sound (Extended Data Fig. 1). Thus, the model covers large part of the auditory processes from the stage before carrier frequency analysis by a cochlea to that making final categorization. This make our model suitable for explaining entire cascade of the auditory system with as little assumptions as possible, unlike in the typical auditory studies which assume frequency-decomposed inputs such as spectrograms. The classification accuracy of the trained DNN was 45.1% (Extended Data Fig.2). We confirmed that depth of the network is necessary to achieve high classification accuracy (Extended Data Fig. 3). Although the classification accuracy was not as good as that reported in other studies^25^, this difference in performance is reasonable when considering that the previous studies used longer (5 s) sound segments for categorization.

### Emerging selectivity to AM frequency

The aim of the present study is to understand the functional significance of the empirically-revealed AM coding scheme in the auditory system, by comparing the AM representation in the trained DNN and that in the auditory system. To enable direct comparison, we simulated experimental approaches of typical neurophysiological studies. Specifically, we conducted “single unit recording” on each unit in the DNN while presenting a sinusoidally amplitude-modulated sound stimulus (Fig. 2a, b). A single unit responded differently to the stimuli with different AM frequencies (Fig. 2c as examples). From the recorded unit activity, we calculated the degree of synchrony to the stimulus AM frequency and the average magnitude of the activity. The synchrony and the average activity as functions of the AM frequency, called a temporal modulation transfer function (tMTF; Fig. 2d, top panel) and a rate modulation transfer function (rMTF; Fig. 2d, bottom panel), characterize tuning to AM frequency in terms of temporal and rate coding, respectively^7^.

**Fig. 2.**
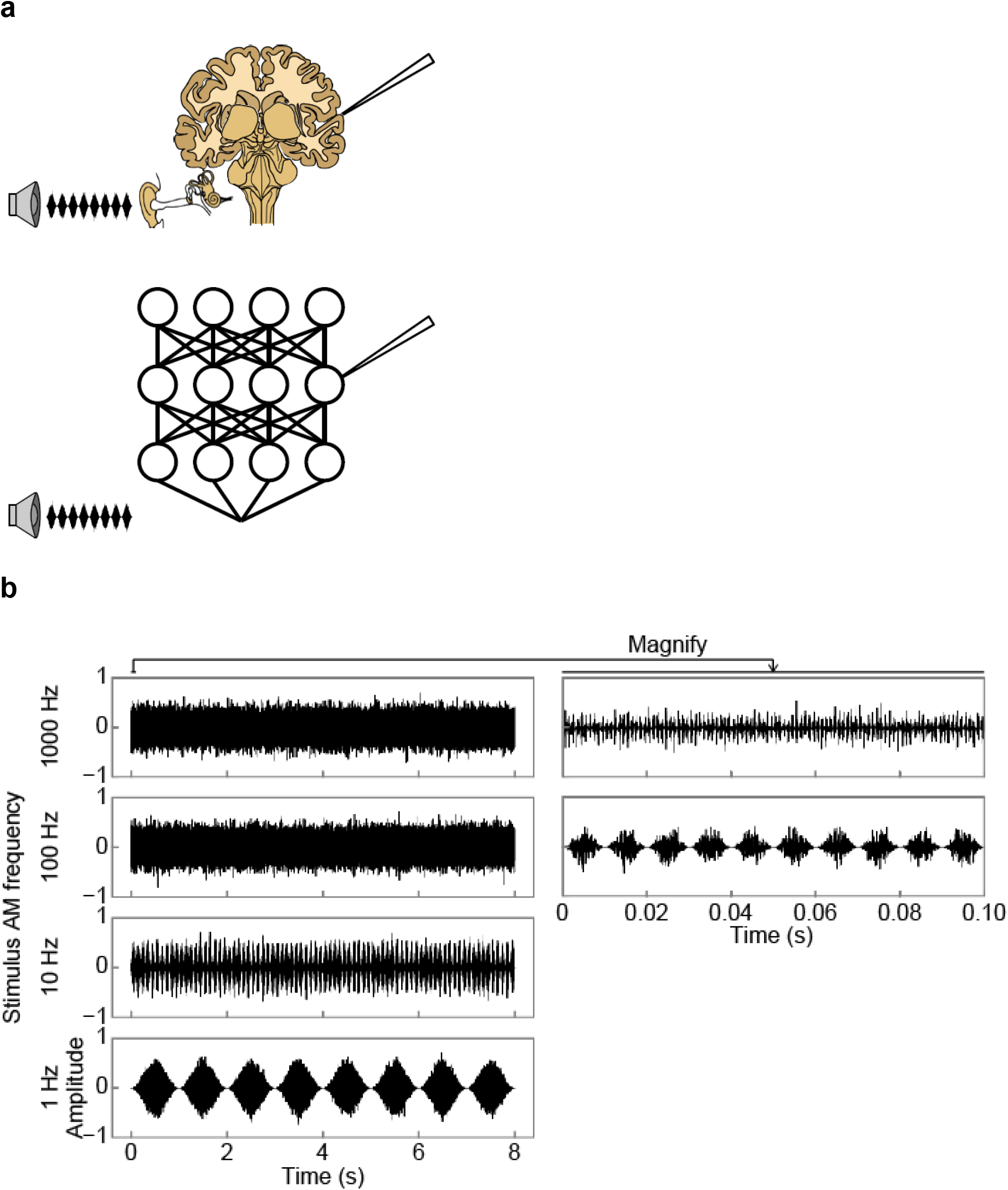
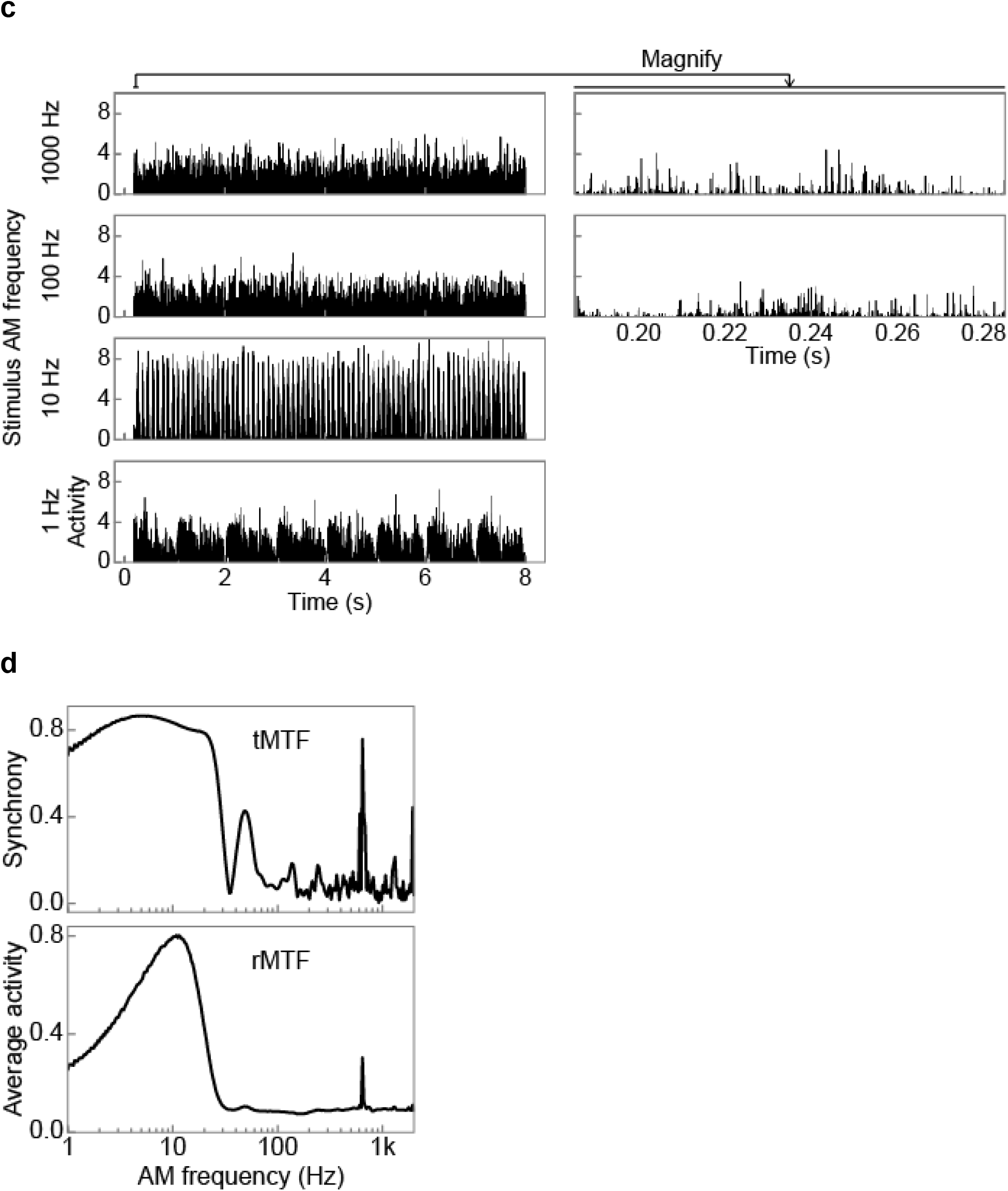
Single unit recording in the DNN. (a) Illustrations of single unit recording in a brain (top) and in a DNN (bottom). In physiological experiments, neural activities are recorded while presenting an AM sound stimulus to the animal. We simulated the method and recorded unit activities of the DNN processing an AM sound stimulus. (b) Examples of AM stimuli with 1, 10, 100, and 1000 Hz AM frequency. The carrier was white noise. (c) Examples of responses to the AM stimuli in (b) in a single unit. A unit in the 8th layer is chosen as an example. Responses to the stimuli with different AM frequencies appeared different. (d) An example of tMTF (top) and rMTF (bottom) in the same unit as (c). A tMTF and an rMTF is defined as synchrony to the stimulus AM frequency and the average activity as functions of AM frequency, respectively. The unit exhibited the low-pass type tMTF and the band-pass type rMTF.

Fig. 3a shows MTFs of representative units in the 1st (i.e., closest to the input), 5th, 9th and 13th (i.e., closest to the output) layers. As in typical physiological experiments, we classified the MTFs into low-pass, band-pass, high-pass or flat types according to certain criteria (see the Methods). Most units exhibited low-pass, band-pass, or flat MTFs, and a negligible number of units exhibited the high-pass type (Fig. 3b). All MTFs in the 1st layer were flat, indicating the 1st layer did not tune to AM frequencies. In the 5th layer, units with low-pass or band-pass tMTFs appeared and a very small number of units with low-pass rMTFs were observed. In the 9th and higher layer, magnitude of tMTF generally increased and the number of units with low-pass or band-pass rMTFs also increased. Heatmaps of all tMTFs normalized by their peaks reveal downward shift of the distribution of the preferred AM frequencies from 5th layer to the highest layer, and distinct tuning in rMTFs appeared from 9th layer and above (Fig. 3c).

**Fig. 3.**
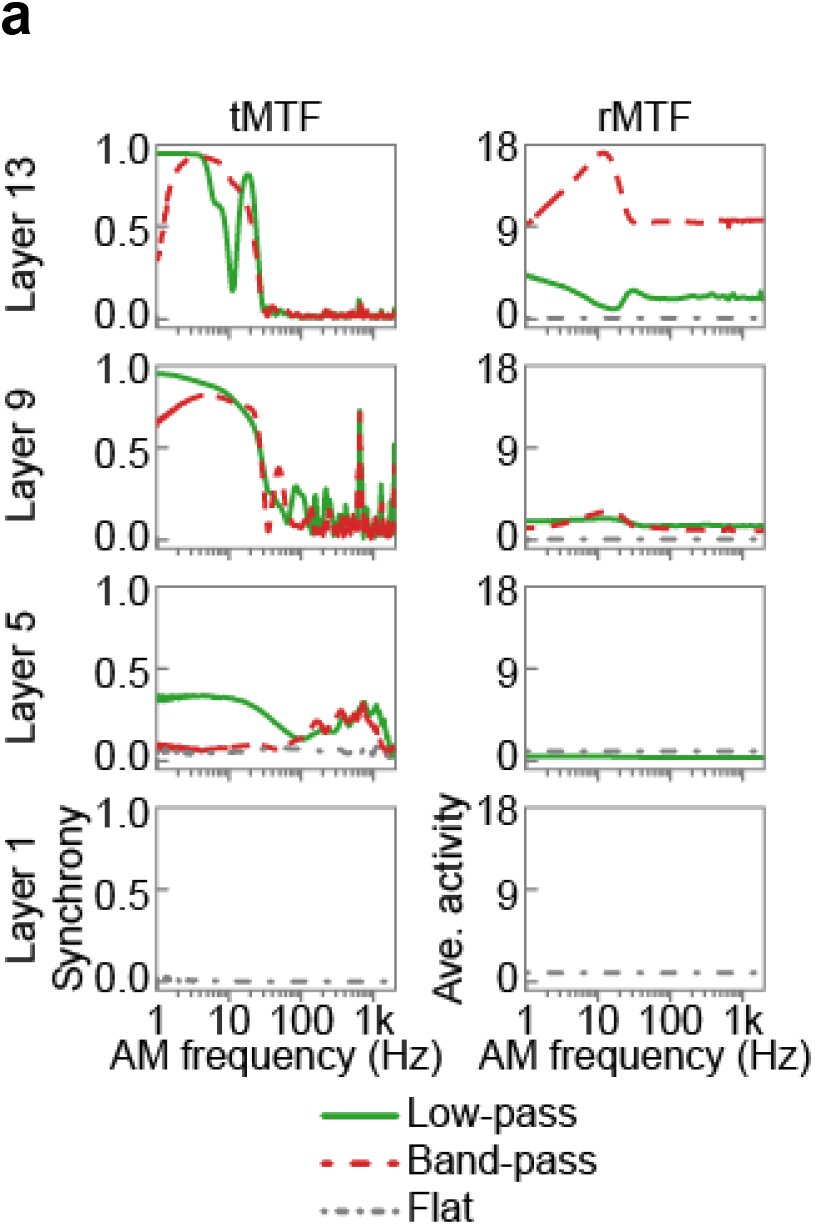
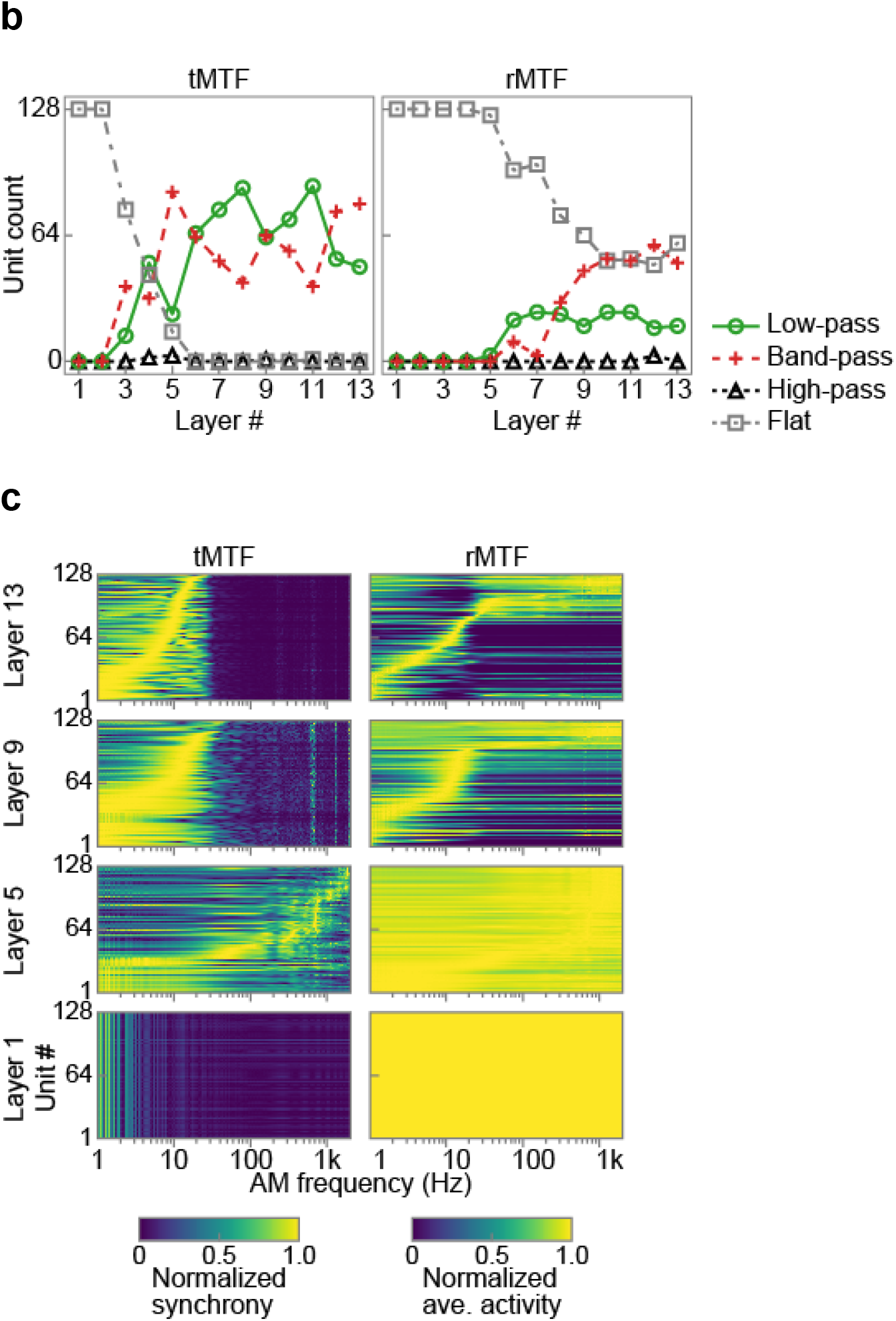
Emergent AM tunings in the DNN. (a) Examples of tMTFs (left panels), and rMTFs (right panels) in layer 1, 5, 9 and 13. The layers are sorted vertically from bottom to top. One example of a low-pass (a solid green line), a band-pass (a dashed red line), and a flat (a dash-dotted grey line) MTF is shown for each layer. In the 1st layer, all MTFs were flat. In the 5th layer significant synchrony to the stimulus AM was observed. In the 9th and 13th layer the synchrony at the lower AM frequencies increased. The magnitude of rate-based responses, shown as the heights of the rMTFs appeared gradually increasing with ascending the layers. (b) The number of units with the low-pass (solid green lines with circles), band-pass (dashed red lines with crosses), high-pass (dotted black lines with triangles), and flat (dash-dotted grey lines with squares) type tMTF (left panel) and rMTF (right panel). Most MTFs were low-pass, band-pass, or flat type. With ascending the layer, the number of low-pass and band-pass MTFs increased. The increase started at higher layer for rate coding than for temporal coding. (c) Heatmaps of all tMTFs (left) and rMTFs (right) in layer 1, 5, 9, and 13. MTFs are normalized by their peak values for better visualization. The units are sorted vertically by their peak AM frequencies. As ascending the layer from the layer 5, the effective AM frequency for inducing synchrony appeared to decrease, and the distinction between darker and brighter area in the rMTFs appeared to become clearer. In some layers, distinct peaks and notches appeared commonly across different units at particular AM frequencies (observed as the vertical lines in tMTFs). We have no clear explanation for this, but this is perhaps due to artefacts of discrete convolutional operation.

### Comparison with the auditory system

As in typical neurophysiological studies, the MTF of a unit was characterized by its best modulation frequency (BMF), the frequency at which the neuron shows the largest synchrony or average activity, and its upper cutoff frequency (UCF), the frequency at which the synchrony or average activity starts to decrease. BMF and UCF of temporal and rate coding are referred to as tBMF/tUCF and rBMF/rUCF, respectively. In the 1st and 2nd layers no BMFs or UCFs were definable since all MTFs were flat (Fig. 4a, b). In the 3rd and 4th layers, units with low tBMFs and tUCFs appeared, but no rBMFs or rUCFs were definable. In the 5th layer, tBMFs and tUCFs tended to be high, and small number units exhibited definable rBMFs and rUCFs. As ascending the layer cascade from the 5th layer, the mode tBMF/tUCF decreased and the number of units with definable rBMFs/rUCFs increased. In sum, the distribution of tBMFs and tUCFs shifted towards lower AM frequencies as ascending from the middle to high layers (Fig. 4a, left panels) and that the units that code AM frequency by their average activities appear only in the higher layers (Fig. 4a, right panels, and Fig. 4b).

**Fig. 4.**
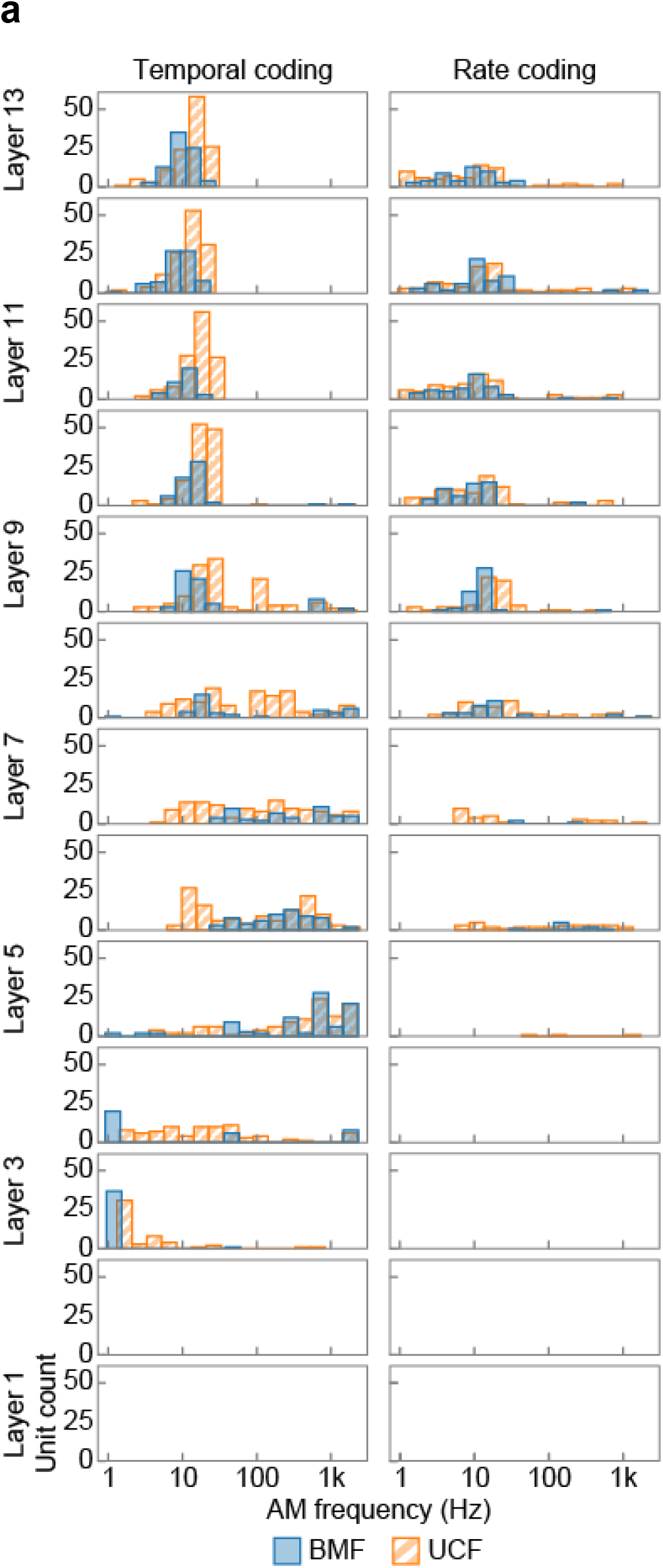
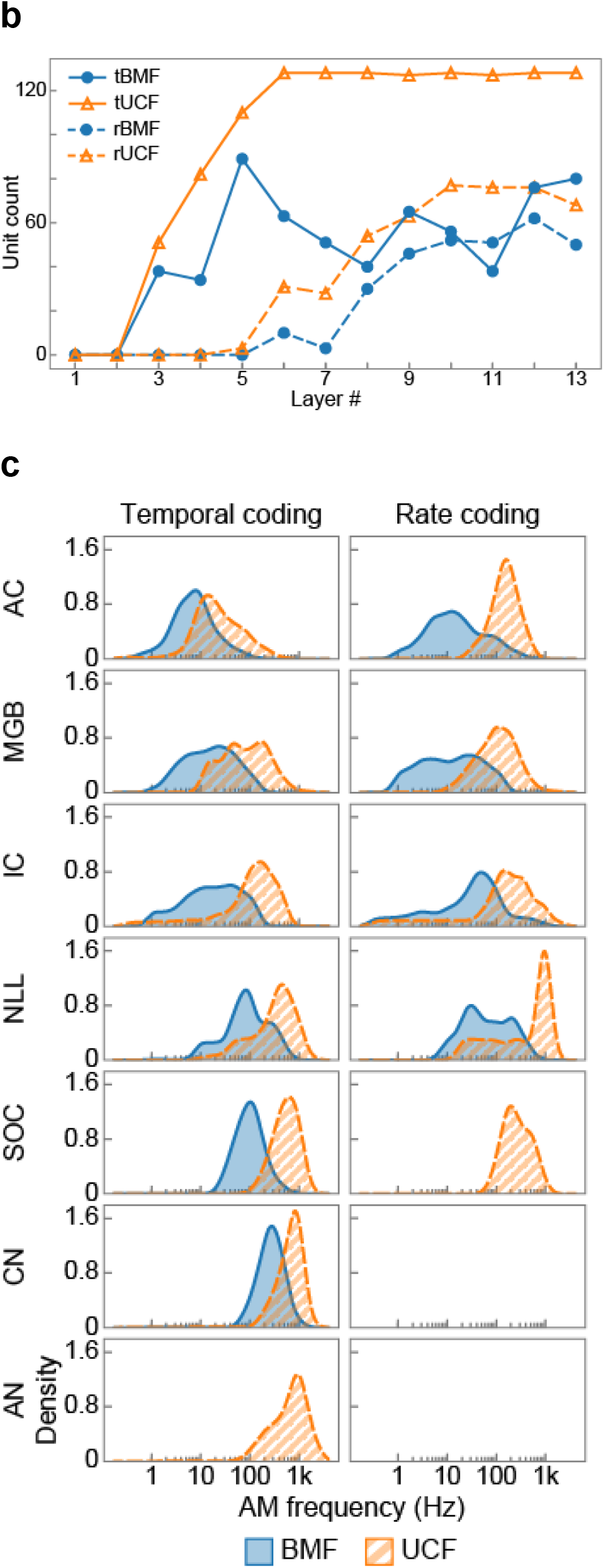
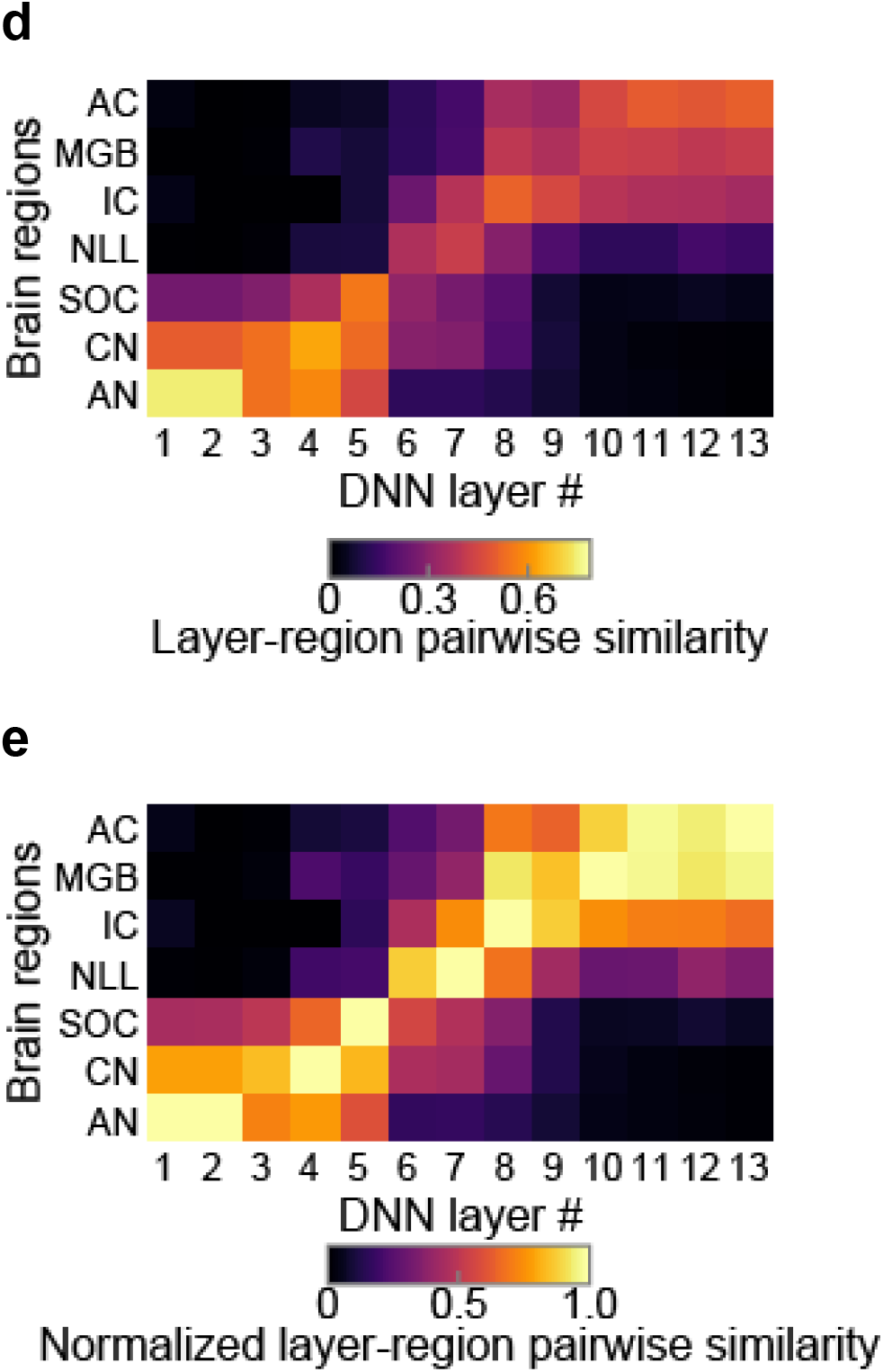
Similarity to the auditory system throughout the entire cascade. (a) Histograms of BMF (filled blue bars) and UCF (hatched orange bars) of temporal (left panels) and rate (right panels) coding in each layer. The layers are sorted vertically from bottom to top. In the 1st and 2nd layer, no units exhibited definable tBMF or tUCF. In the 3rd and 4th layer, the tBMFs and tUCFs covered wide range of the AM frequency, majority of them being low. As ascending from 5th layer, the tBMFs and tUCFs appeared to decrease. As for rate coding, in the 1st to 4th layers, no units exhibited definable rBMF or rUCF. In the 5th layer small number of high rBMFs and rUCFs appeared. As ascending from the 5th layer, the number of units with definable tBMFs and tUCFs increased. (b) The number of units with definable BMF (filled blue circles) and UCF (open orange triangles) of temporal (solid lines) and rate (dashed lines) coding. As ascending the layers, the number of definable units increased. The number of units with definable rBMF and rUCF started increasing in higher layers than those with definable tBMF and tUCF. In other words, rate coding are performed in higher layers than temporal coding. (c) Distributions of BMF (filled blue areas) and UCF (hatched orange areas) of temporal (left panels) and rate (right panels) coding in each region in the auditory system. Regions are sorted vertically from the peripheral regions (bottom panels) to the central (top panels). No distribution of tBMF is reported in AN. The tBMFs and tUCFs gradually decrease from the periphery to the central. No distribution is reported for rate coding in the peripheral regions probably because peripheral regions do not code AM frequency by the spike rate. (d) Layer-region pairwise similarity of the AM representation in the DNN layers (horizontal axis) and that in the regions in the auditory system (vertical axis). Pairs of layers and regions with large similarity appeared in diagonal. (e) Layer-region pairwise similarity normalized by the maximum value of each brain region. The diagonal pairs with large similarity are more clearly observed.

The patterns of the BMF/UCF distributions reminds us of the well-known characteristics of the auditory pathway, i.e., decrease of synchronizing AM frequency^5,7^ and time-to-rate conversion of AM coding^7^. Fig. 4c visualizes the distributions of BMFs and UCFs in the auditory system, combining previously reported distributions in each of the 7 brain regions: auditory nerves (AN)^26,27^, cochlear nucleus (CN)^26,28–31^, the superior olivary complex (SOC)^29,32^, the nuclei of the lateral lemniscus (NLL)^33–35^, the inferior colliculus (IC)^36–40^, the medial geniculate body (MGB)^41–43^, and the auditory cortex (AC)^40,44–51^. In the peripheral regions tBMFs and tUCFs clustered around high AM frequencies, and as ascending towards the central, the mode frequencies decreased. RBMFs are only reported in NLL or above, and rUCFs are in SOC or above.

The meta-analysis of the neurophysiological studies suggests qualitative similarity of the distribution of the BMF and UCF in the DNN and those in the auditory system. Next, we quantitatively compared those distributions. For each of the tBMF, tUCF, rBMF, and rUCF, we calculated the similarity between the distribution in each layer of the DNN and the distribution in each region in the auditory system (Extended Data Fig. 4), and averaged them to yield the layer-region pairwise similarity (Fig. 4d). Pairs of the DNN layer and the brain region with large similarity appeared in the diagonal direction, indicating that lower, middle, higher DNN layers are similar to peripheral, middle, and central brain regions, respectively. This lower-periphery, middle-middle, and higher-central similarity is more clearly observed if we normalized the pairwise similarity by the maximum in each brain region (Fig. 4e).

### Relationship to optimization

Is the observed similarity of the entire cascade between the DNN and the auditory system due to the convolutional architecture inherent to the DNN^52^ or the consequence of optimization of the filter weights and biases for the classification task? To test these possibilities, we measured MTFs in the DNN before and during the optimization. Before the optimization, no unit showed clear selectivity to AM frequency, and there appeared little transformation of MTFs across layers (Extended Data Fig. 5, left panel). All layers were similar to the peripheral regions (Fig. 5a).

**Fig. 5.**
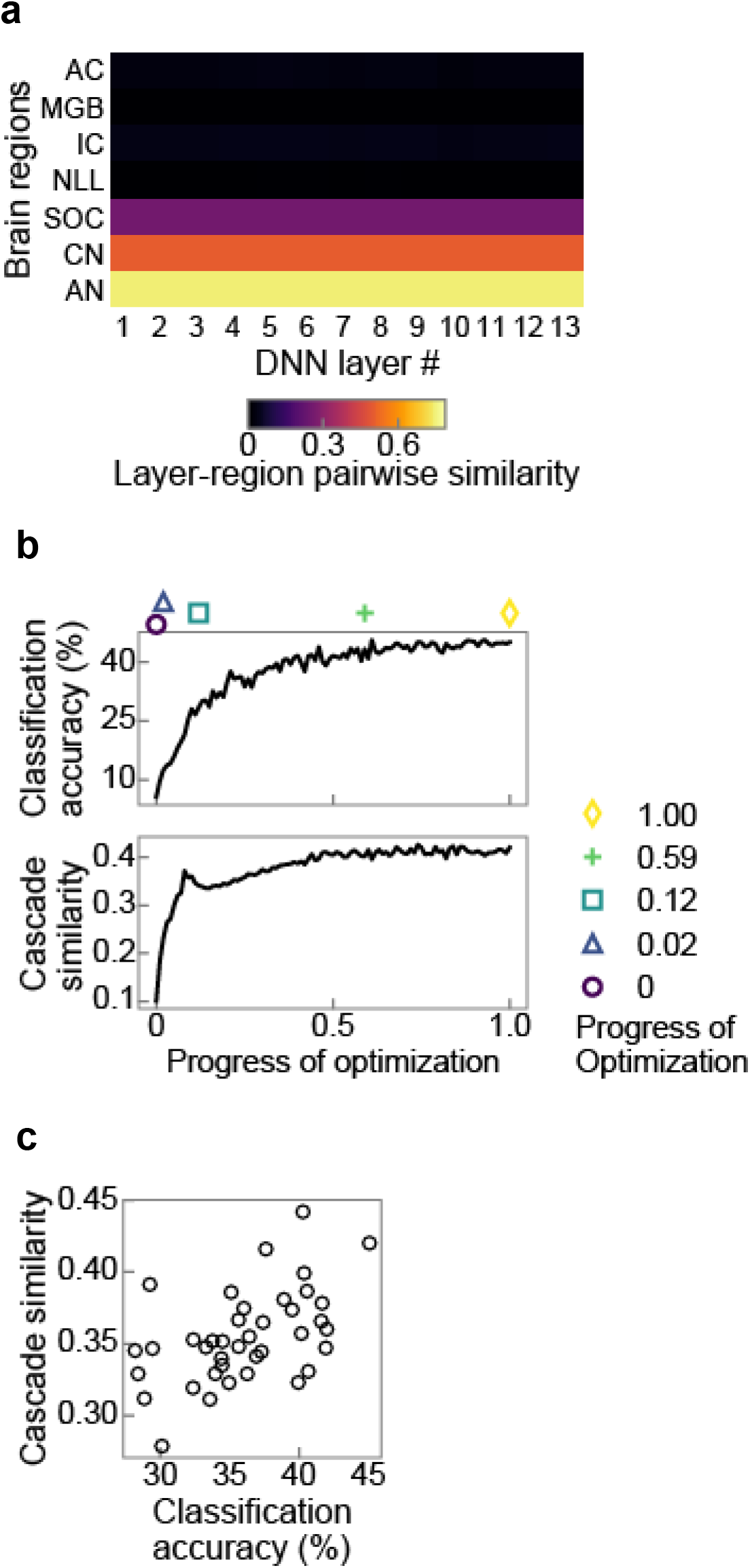
Similarity correlated with classification accuracy. (a) Pairwise similarity of the DNN before optimization. Other conventions are the same as in Fig. 4d. All layers in the DNN were similar to the peripheral regions. (b) The classification accuracy (top) and the similarity of the entire cascade (bottom) as functions of the progress of optimization. The progress of optimization, shown in the horizontal axis, is linearly normalized so that the value takes 1 at the end of the optimization. The classification accuracy and the similarity increased as the optimization progressed, indicating the emergence of the auditory-system-like AM coding during the optimization. Coloured markers indicates the points at which layer-wise similarities were calculated in Fig. 6a. (c) The similarities of the DNNs with various architectures, plotted against their classification accuracies. The correlation indicates that AM representation in the better-performing DNNs are more similar to the auditory system.

As the optimization progressed, classification accuracy increased as expected (Fig. 5b, top panel). In parallel, auditory-system-like AM tuning gradually emerged (Extended Data Fig. 5). We evaluated the similarity over the entire cascades by measuring the degree of diagonality of the pairwise similarity matrix (Extended Data Fig. 6), and defined it as the cascade similarity. A greater value of the cascade similarity indicates that, in the pairwise similarity matrix, cells around the diagonal line exhibit large similarity and cells around left-top and right-bottom corners exhibit small similarity. The cascade similarity increased as the optimization progressed (Fig. 5b, bottom panel), and correlated with the classification accuracy very well (Spearman’s rank correlation coefficient *ρ* = 0.84, *p* = 8.57×10^−28^). The results indicate that the AM representation in the DNN emerged during the optimization.

The above results indicate that similarity to the auditory system, as well as classification accuracy, depends on the parameters of the DNN. Generally, classification accuracy of a DNN also depends on its architecture^53,54^, and cascade similarity, too. We trained DNNs with various other architectures and examined them with the same physiological analysis. The classification accuracy of those DNNs varied between 28.2% and 45.1%. The patterns of the layer-region pairwise similarity also varied among the architectures (Extended Data Fig. 7), and the cascade similarity correlated with the classification accuracy (Fig. 5c; Spearman’s rank correlation coefficient *ρ* = 0.51, *p* = 8.08×10^−4^). The results indicate that AM representation in better-performing DNNs are more similar to that in the auditory system. Taken together, similarity to the auditory system correlated with classification accuracy both across different model parameters and across different architectures, suggesting strong relationship of the auditory AM representation to parameter optimization, but not to the convolutional operation alone.

### Different factors for different regions

The changing pattern of the layer-region pairwise similarity during optimization indicates that auditory-system-like AM tuning first emerged in the upper layers, followed by middle layers (Extended Data Fig. 5). This pattern is more clearly seen when we calculated the similarity to the auditory system in each layer, which we call layer-wise similarity (Fig. 6a, Extended Data Fig. 6). Before optimization, AM representation was similar to the auditory system only in the lower layers. As optimization progressed, similarity in the upper layers rapidly increased, and then similarity in the middle layers increased. The result implies that multiple factors can underlie these across-layer differences in the evolution patterns. To isolate the possible factors in each region, we conducted the following four control experiments, expecting to see different degrees of similarity emerges in different layers depending on the control conditions.

**Fig. 6.**
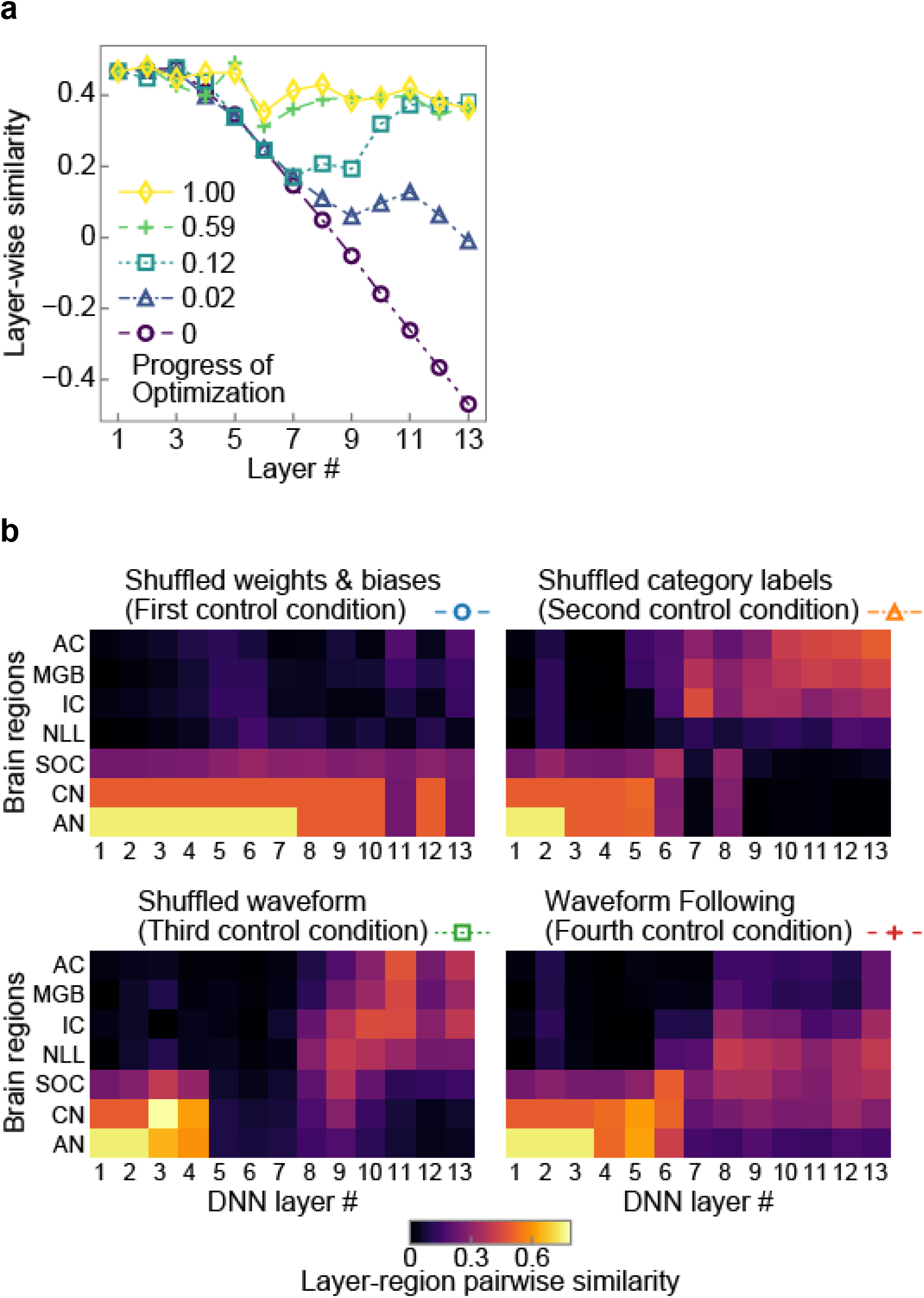
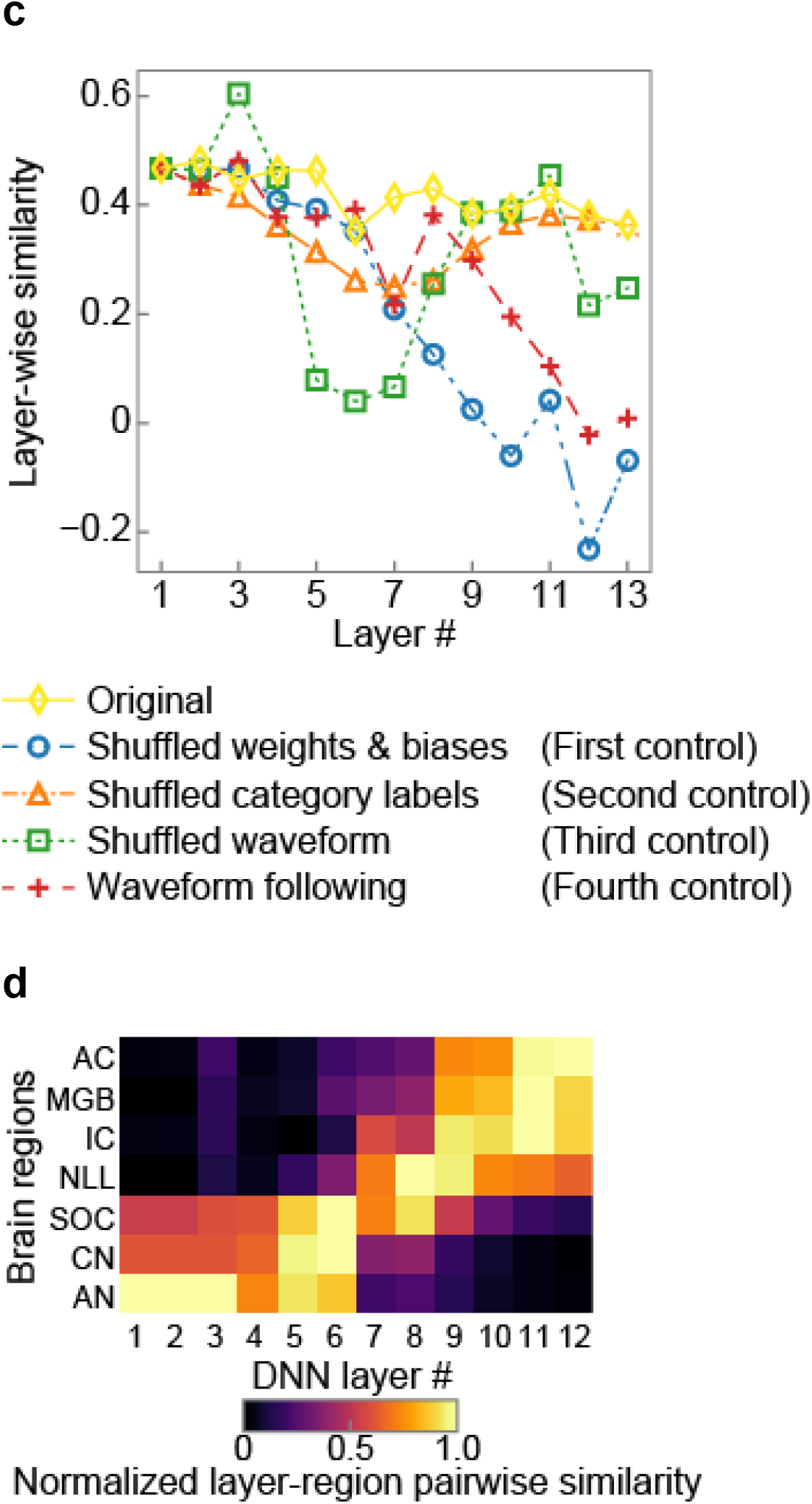
Different factors for different regions and consistency across datasets. (a) Layer-wise similarity at the four intermediate snapshot instances during optimization. Colors, markers, and lines indicate the progress of optimization as indicated by the legend and in Fig. 5b. As optimization progresses, similarity in the higher layers rapidly increased, followed by the middle layers. (b) Pairwise similarity in the control experiments. Coloured markers and lines by the panel titles indicate the types of the control conditions as in (c). Other conventions are the same as in Fig. 4d. (c) Layer-wise similarity in the control experiments. The similarities in the original condition (yellow diamonds and solid line) are also shown. The lower layers were similar to the peripheral regions in all conditions. The middle layers were simlar to the middle regions only in the original and fourth conditions, and the higher layers were similar to the central regions only in the original, second, and third conditions. (d) Layer-region pairwise similarity of the DNN trained on a speech dataset. Other conventions are the same as in Fig. 4e. The lower layers are similar to the peripheral regions and the higher layers are similar to the central regions, indicating auditory-system-like AM representation consistently emerged from the speech dataset.

First, we tested the effect of specific assignment of the parameters. Our DNN has two types of trainable parameters: filter weights and biases. Examination of the parameters of the optimized and preoptimized DNN reveals that the distribution of the bias values in the optimized DNN deviated largely from 0, the initial fixed values before optimization, although the distribution of the filter weights did not change very much (Extended Data Fig. 8). It is possible, for example, that overall changes in the bias values that take place in some layers had an effect to amplify or suppress the higher representation. We tested this possibility by randomly shuffling the filter weights and biases within each layer. The resulting AM representation in all layers were similar to that in the peripheral regions in the auditory system (Fig. 6b, left top panel). A few units in the upper layers appeared to exhibit some tuning to low AM frequency, but majority of the units did not show significant AM tuning (Extended Data Fig. 9, left column). Thus, the result disproved the effect of overall distribution of the parameters, and confirmed the importance of the specific assignment of the parameters for auditory-system-like AM tuning.

The second and third control experiments tested the effect of data structure. It has been shown that a DNN is capable of learning the input-output correspondence even by training on data with random category labels or data without natural statistics^55^. It can be argued that the process of optimization, but not the data structure, is the essential factor for inducing AM tuning. To test this possibility, we trained the DNN with unnatural data. In the second control condition, the input-output correspondence was destroyed by shuffling category labels, making accurate classification of novel data impossible. In the third control condition, the structure of the input waveform was destroyed by shuffling waveform in each sound. The DNN was able to classify the novel sounds with some accuracy probably because the waveform shuffled within each sound retained its overall amplitude distribution, although both frequency and temporal statistics are completely destroyed. The trained DNNs in these two conditions exhibited auditory-system-like AM representation only in the lower and upper layers, but the middle layers failed to exhibit AM representation similar to the middle auditory regions (Fig. 6b, right top and left bottom panels, Fig. 6c, orange triangles and green squares, Extended Data Fig. 9, second and third columns). When trained on shuffled labels, very few units in the middle layers appeared to exhibit AM tuning. When trained on shuffled waveform, units in the middle layers appeared to exhibit some AM-frequency tuning but they synchronized to much higher AM frequency than neurons in the auditory system, making relatively higher layers around layers 8-10 resemble relatively peripheral regions such as CN and SOC. The results indicate that mid-level AM representation requires natural data structure, although that low-level and high-level representation could emerge just by optimizing even to unnatural data.

Finally, the fourth control experiment examined the effect of the optimization objective. A DNN may be optimized not only for an ethologically relevant objective such as sound classification, but also for unnatural objective such as the waveform following task. To test the effect of optimization objective on emerging AM representation, we trained the DNN for the waveform following task. Specifically, the DNN was trained to copy the input waveform (Extended Data Fig. 10). This task has no biological significance and is trivial in the sense of signal processing. A successful network should maintain information of the input waveform throughout the depth of layers with non-linear processes without lowpass filtering. The AM representation in middle to upper layer was to some degree similar to the middle brain regions, but no layers exhibited AM representation similar to the central brain regions (Fig. 6b, right bottom panel, Fig. 6c, red crosses, Extended Data Fig. 9, right column). In the higher layers, MTFs did not show clear tuning, and the tBMFs and tUCFs were higher than the central auditory regions, making the higher layers resembling middle auditory regions. The result indicates that emergence of auditory-system-like AM tuning in the higher layers requires natural objectives, and the waveform following task did not induce such representation even if the input data were the natural sounds.

Taken together, modification of the weight and bias assignment, the category labels, the sound statistics, and the optimization objective deteriorated the auditory-system-like AM representation in some layers. Lower layers never exhibit AM tuning probably because of the nature of the cascading architecture. The middle layers exhibited auditory-system-like AM tuning when trained on the natural input sounds and the proper sound-category correspondence. The upper layers exhibited auditory-system-like AM tuning when optimized for the categorization task but not for the waveform following task (Table 1).

**Table 1.** Major factors for AM representation in different regions.

| Regions | Major factor |
| --- | --- |
| Lower | Cascading architecture |
| Middle | Data naturalness |
| Higher | Optimization objective |

### Generality across datasets

It can be argued whether the obtained results were specific to our choice of the dataset, animal vocalizations and environmental sounds. Previous studies show positive pieces of evidence for the generality across datasets. A DNN trained on one dataset can be transferred to another task with only small modification^56^. Also, an efficient-code model trained for substantially different sound datasets, one consisting of human speech and the other of animal vocalizations and environmental sounds, exhibits quantitatively similar representation of carrier frequency^57^. To test the generality of the finding of the present study across datasets, we conducted the “physiology” in a DNN optimized for phoneme classification of speech sounds. A segment of speech sounds in the dataset was labelled with corresponding phoneme, an element of vocalization in speech.

The DNN trained on the speech derived essentially the same conclusions as those shown by the DNN for the animal and environmental sounds. The layer-region pairwise similarity matrix exhibited the diagonal pattern (Fig. 6d): Lower layers were similar to peripheral regions, middle layers to middle regions, and higher layers to central regions. The similarity emerged during the optimization, and was weak in the control conditions (Extended Data Fig. 11a, b). The similarities in the DNNs with various architectures correlated with the classification accuracy (Extended Data Fig. 11c; Spearman’s rank correlation coefficient *ρ* = 0.33, *p* = 3.91×10^−2^).

### Tuning to carrier frequency

Other than tuning to AM frequency, one of the frequently measured characteristics of auditory neuron is tuning to carrier frequency^58,59^. We calculated temporal average of the activities in each unit in response to a sinusoid with various frequencies and amplitudes (Extended Data Fig. 12a). The responses generally increased as the amplitude of the input increased, but some units in higher layers showed non-monotonic responses to the input amplitude. For instance, in the layer 13, the unit shown in the right panel in Extended Data Fig. 12a exhibited large responses to ~ 30 dB, 400 Hz tone, but the response was smaller to the tone with larger amplitude. As in the neurophysiological studies, a unit was characterized by a frequency tuning curve, the minimum stimulus amplitude which gives larger response than a certain threshold (Extended Data Fig. 12a, grey lines, Extended Data Fig. 12b). Frequency tuning curves in the lower (1st to 3rd) layers appeared to exhibit many peaks. Those in the middle layers (around 5th layer) appeared to exhibit single large peaks and multiple small peaks. The large peaks appeared to span wide range of the carrier frequency as a population (Extended Data Fig. 12b), which may be interpreted as a band-pass filter bank. Frequency tuning curves in the higher (8th to 13th) layers appeared to be more complex without clear bandpass-like tunings even as a population. The results were in contrast to the auditory system. Neurons usually exhibit frequency tuning with a sharp single peak, which is likely to originate from frequency decomposition performed in the cochlea. We did not explicitly conducted spectral decomposition of the input sound but directly fed raw waveforms to the DNN. The results suggest that frequency decomposition in the cochlea may be essential for auditory-system-like carrier frequency tuning but not for auditory-system-like AM tuning.

## Discussion

We found that a DNN optimized for sound classification exhibits AM representation similar to the auditory system throughout the entire cascade of the signal processing. The lower layers in the DNN were similar to the peripheral regions, the middle layers to the middle regions, and the higher layers to the central regions. Such representation gradually emerged during the optimization and correlated with the classification accuracy. The control experiments suggest that essential factors for AM representation in the lower layers, middle layers, and higher layers are the cascading architecture, data naturalness, and optimization objectives, respectively. Such representation was consistently observed in the DNNs trained on different datasets. The similarity of the entire cascade was demonstrated because our DNN performs sound recognition from a raw sound waveform. Since our DNN was not designed or trained to reproduce any physiological or anatomical properties of the auditory system including cochlear frequency decomposition, the results should reflect only the nature of the task and the data. It would be an important finding that the characteristics regarding AM coding, which are essential for auditory perception, are common in a DNN and the auditory system. These results suggest that AM representation in the auditory system might also be the consequence of optimizing to the sound recognition in the real world, which could emerge during evolution and development.

### AM Representation in the lower, middle, and higher layers

Our results suggest that AM representation in the lower layers is due to the cascading nature of the system. A DNN performs highly nonlinear operation by cascading close-to-linear operations. Perhaps this is also what happens in the auditory system. Neurons in each layer performs relatively simple operation, which may lead to little sensitivity to AM frequency in the peripheral regions.

The representation in the middle and higher layers, however, depended on the optimization condition. The representation in the middle layers were similar to that of the auditory system only in the DNN with high classification accuracy, but not in the DNNs with poor classification accuracy, the DNN halfway in the optimization process, or the DNN trained with unrealistic data. This suggests that mid-level AM representation is essential for effective representation of natural sounds. On the other hand, AM representation in the higher layers were similar to the auditory system in all of these conditions but the waveform following task. This suggest that task natures are determinant factors for forming high-level AM representation, perhaps because higher representation is more directly used for final decision than middle or lower representation. In other words, whatever the lower representation is, the role of the higher layers are to derive appropriate outputs for the specific task from the lower representation

### Decreasing temporal resolution for sound classification

Both of the two prominent characteristics of the auditory AM coding, decrease of synchronizing AM frequency and time-to-rate conversion, involve decrease of temporal resolution of the transmitted signals. The above discussion regarding representation in higher layers suggests that encoding information of sound categories with low temporal resolution may be beneficial for classification tasks.

The next question is why such coding scheme is beneficial. The following discussion might explain the reason. In our setting, as in the typical classification task with a DNN, the larger the value in each unit in the classification layer (the layer above the 13th layer), the larger the score will be for the particular category. The final output category is the one assigned to the unit with the maximum value. If the units synchronize to the amplitude envelope of the input sound, which wax and wane with time, the output category would be temporally unstable. On the other hand, if the activity of an output unit is large all the time, the score for the category will be kept large. The latter case would be more preferable for classification tasks.

In the real world, recognizing the stimulus category would be more important than synchronizing to the stimulus, and animals might be better at sound classification than synchronizing to the sound. This notion is supported by the well-known phenomenon that in a synchronization tapping task humans tend to respond slightly earlier than the correct timing^60^, suggesting that we tap according to the internally generated rhythm but not react after hearing the ongoing sound. Other animals which have the ability to act synchronously to a stimulus exhibit similar behaviour^61^. These animals (including humans) might first recognize the frequency of the stimulus envelope and then generate rhythm at the recognized AM frequency. Such behaviour might also be observed if a DNN optimized for sound classification is forced to perform a synchronization tapping task.

A reader who is familiar with a convolutional DNN may think that low temporal resolution in the higher layer is trivial if each layer performs pooling operation, which temporally downsamples the input waveform. However, this is not the case for our DNN, in which no pooling was performed. Thus layers in our DNN does not necessarily downsample the input. Indeed the DNN trained for the waveform following task did not decrease temporal resolution very much.

### Frequency tuning

Our DNN did not exhibit sharp single peaks in the frequency tuning curves as widely found in the auditory system, while some studies report auditory-like frequency tuning emerging in a DNN with different architecture from ours^64,65^. In the auditory system, frequency tuning of a neuron is largely affected by mechanical and physical properties of the cochlea^59^. Although investigating what determines the shape of a frequency tuning curve in a DNN is beyond the scope of this study, some architectural constraints might be necessary for inducing similarity to the auditory system in the carrier frequency domain.

Several other modelling works try to explain AM coding in the auditory system with anatomical and physiological assumption including frequency decomposition in a cochlea^23,66,67^. A message brought from the present study, which did not incorporate cochlear frequency decomposition, is that sharp frequency tuning may not be necessary for effective AM representation for natural sound recognition.

### Physiology in a DNN

Our results suggest the effectiveness of analysing computational model using physiological methods. To date various methods have been proposed for analysing representation in a DNN^63^. Most of them rely on differentiability of the DNN, using backpropagation to estimate the optimal input for each unit assuming such an input exists. On the contrary, there is a long history of developing physiological method to elucidate brain functions. Physiologists rely on parametric search over the stimulus space, since backpropagation cannot be applied to the biological neurons^5^. One advantage of our method is that the results are directly comparable with the ones reported in the physiology experiments. By taking advantage of the previously-conducted vast number of neurophysiological studies, we could show the relationship between layers in the DNN and the regions in the entire cascade of the auditory system. Although DNNs have been used to explain sensory representation in several modalities^20–24^, to the best of our knowledge this is the first report of similarity throughout the entire cascade of the sensory processing. The success of our method indicates the future possibility of applying well-established physiological paradigms to explore the functions and mechanisms of a DNN and other complex machine learning models.

From a physiological perspective, this study implies that a DNN may become a useful tool for testing a new hypothesis. Although this study focused on representation of sound envelopes, for which large amount of physiological data are already available, any domain of stimulus parameters can be explored in the same paradigm as ours. As long as the model takes raw data as in this study, physiologists can test their hypothesis on any sensory domains with any kinds of stimuli with much lower costs than actually conducting a pilot physiology experiment.

## Methods

### Task

The task of the DNN was sound classification. Specifically, the task was to estimate the sound category at the last timeframe of a sound with certain duration (0.19 s for natural sounds and 0.26 s for speech). A classification accuracy is defined as an average of the correct classification rate for each category, which is the number of timeframes correctly estimated as the particular category divided by the number of total timeframes of the category.

### Dataset

The following two datasets were used to train DNNs. The first one consists of non-human natural sound, which is a subset of ESC-50^68^. The original dataset contains 50 sound categories with 40 sounds for each category. From the original dataset we used 18 categories which are not produced by human activities. Each entry in the original dataset contains a sound waveform of length less than 5 s and the category of the sound. In this study we excluded silent intervals, resulting in the total length of 53.9 minutes. The original dataset is divided in 5 folds for cross validation. We used fold #5 for validation and the other fold for training. The sound format was 44.1 kHz 16 bit linear PCM.

The second dataset consists of speech sound^69^. Each entry in the dataset contains a sound waveform of a single spoken sentence, phoneme categories, and time intervals of each phoneme. The original number of phoneme categories is 61. We merged some categories in accordance with the previous study^70,71^, resulting in 39 categories. The average duration and the total duration of the sound is 3.1 s and 3.3 hours, respectively. The data is originally divided in validation set and training set. In this study we followed the original division. The validation set and training set contains speech of 24 and 462 speakers, respectively. The speakers and the sentences in the two dataset did not overlap. The sound format was 16 kHz 16 bit linear PCM.

### Network architecture

Our DNN consisted of a stack of dilated convolutional layers^62^ (Extended Data Fig. 1), in which convolutional filters are evenly dilated in time. Convolution is conducted along the time axis. Each layer performs dilated convolution to the output of the previous layer and applies rectification as an activation function. The activation function was an exponential linear unit^72^. The first layer directly took samples of raw waveforms as an input. Each layer contains multiple units. In our setting, each layer contains same number of units for simplicity. The units in the highest layer is connected to the classification layer without convolution. The number of the units in the classification layer was the number of the categories. The classification layer was omitted from the physiological analysis.

We used DNNs with 13 layers, each containing 128 units, for non-human sound, and DNNs with 12 layers, each containing 64 units, for speech. The number of layers and the number of units in each layer were determined based on the pilot study and fixed to the value throughout the study. In the pilot study DNNs with various number of layers and units were trained using random portion of the training set. The filter length was 2, and the dilation length was 2 to the power of the layer number^62^. The number of layers and the number of units in each layer that gave the best classification accuracy on the other portion of the training set were used in the following study.

We tested multiple architectures with random filter and dilation length in each convolutional layer and selected the DNN which achieved the best classification accuracy on the novel dataset (Extended Data Table 1). The filter size and dilation length was randomly chosen for each layer with constraints that the filter size does not exceed 8 and the total input length for the whole DNN, which is equal to the length of the input time window of the highest layer, does not exceed 8192 (~ 0.19 s) for non-human sound and 4096 (~ 0.26 s) for speech. The number of layers and the number of units in each layer were fixed as mentioned in the previous paragraph.

### Optimization

The DNNs were trained on the training set, and the classification accuracy were calculated on the validation set. The initial filter weights were randomly sampled and biases were set to 0 in accordance with the previous study^73^. The filter weights and biases were updated using Eve algorithm^74^ with softmax cross entropy as the cost function. The number of iteration for parameter update was determined to the value which gave the best classification accuracy on random portion of the training set trained on the other portion of the training set.

### Physiological analysis of a DNN

For physiological analysis of a DNN a sound stimulus was fed to the DNN and the values of each unit were recorded. The root mean square (RMS) of the input sound was adjusted to the mean RMS of the training set. Before analysis, 1 was added to the values of all units because the minimum possible value of the activation function is –1^72^.

The stimulus was 8 s of sinusoidally amplitude modulated white noise (Fig. 2b). In the physiological studies tuning to AM frequency is measured with sinusoidally amplitude-modulated tones with carriers at the neurons’ best frequencies, sinusoidally amplitude-modulated white noises, or click trains. We did not use tones as carriers because many units showed multiple peaks in the tuning curves to carrier frequency or non-monotonic responses to the input amplitude (Extended Data Fig. 12), making it difficult to define the best carrier frequencies.

From the values of each unit the synchrony to the stimulus and the average activity was calculated. The synchrony to the stimulus was quantified by a vector strength^75^. When dealing with spike timing data recorded in biological neurons, each spike is represented as a unit vector with its angle corresponding to the modulator phase at that time, and the vector strength is defined as the length of the average of these unit vectors. Equivalent operations were applied to the continuous output of the DNN unit to derive a value of vector strength (equation 1). The vector strength takes a value between 0, indicating no synchrony, and 1, indicating perfect synchrony.

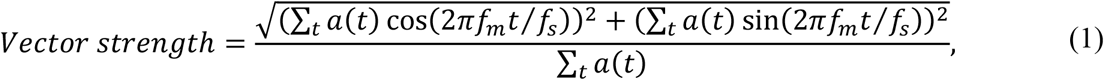

where *t* is an index of the timeframe, *a*(*t*) is the unit activation, *f_s_* is the sampling rate, and *f_m_* is the stimulus AM frequency. The average activity was defined as the temporal average of the values, which could be considered as the DNN version of an average spike rate. The synchrony and the average activity was averaged for 16 instances of the carrier white noise to reduce the effect of stimulus variability. A tMTF and an rMTF was defined as the synchrony and average activity as functions of AM frequency, respectively. In physiology an MTF is usually defined only at the frequencies at which the unit shows statistically significant synchrony or spike rate. Since a statistical test on the results of deterministic model such as our DNN does not make sense, we considered the synchrony or average activities less than a certain threshold as “non-significant” and excluded them from the following analysis. The threshold was arbitrarily set to 0.01 for the synchrony and to 0.01 above the average activity in response to unmodulated white noise for the average activity.

An MTF was classified into one of the following 4 types: low-pass, high-pass, band-pass, or flat. The low-pass type MTF was defined as the one not having values smaller than 80% of its maximum in the frequencies smaller than the peak frequency. The high-pass type MTF were defined as the one not having values smaller than 80% of its maximum in the frequencies larger than the peak frequency. The flat MTF was defined as the one not having values smaller than 80% of its maximum or the one with the peak to peak range less than 0.1. The band-pass MTF was defined as otherwise.

BMFs were calculated from the band-pass type MTFs, and UCFs were calculated from the low-pass and the band-pass type MTFs. BMFs of low-pass, high-pass, or flat MTFs and UCFs of high-pass or flat MTFs were considered as indefinable. The BMF was defined as the modulation frequency at the peak of the MTF. If multiple peaks with the same height exist, the geometric mean of the frequencies was taken. The UCF was calculated in two different ways: one for qualitative visualization in Fig. 4a and the other for quantitative comparisons with specific physiological data of neurons in the literature. The UCF for visualization was defined as the frequency at which the value of the MTF crosses 80% of its maximum. If a MTF had multiple such frequencies, the geometric mean of the frequencies was used. The threshold of the UCF for quantitative comparison with the auditory system varied according to the reference physiology study. They were 50%^35,49^, 80%^26,32^, and 70% (–3 dB)^27–29^ of the maximum, 90%:10% interior division of its minimum and maximum^36^, absolute value of 0.1^26,31^, and the highest frequency that gives significant responses^32,33,36,38,42–44,47,50^. If at no frequency did the MTF cross the threshold, the UCF was considered as indefinable.

Stimuli for calculating a tuning to carrier frequency were tones with various frequencies and amplitudes. The values of each unit was temporally averaged to obtain the response to the particular stimulus. The tuning curve was defined for each frequency as the smallest amplitude inducing the response larger than a certain threshold. In physiological studies thresholds are usually determined arbitrarily. In Extended Data Fig. 12 tuning curves with the thresholds of 0.001, 0.01, and 0.1 are shown.

### Comparison with the auditory system

We extracted the distributions of BMF and UCF reported in the previous physiological studies by digitizing the printed figures in each paper. If multiple figures were available, we chose the clearest figure or the one with most number of neurons. The extracted values were used in qualitative visualization in Fig. 4c and quantitative comparison with the DNNs. For visualization the distributions of all sub-regions and all neuron types in each region in each paper were averaged. Then the distributions of all papers were averaged for each region. The resulting distributions were smoothed with a Gaussian filter with width 0.136 in the logarithmic scale of base 10. For quantitative comparison with a DNN, the similarity of each extracted distribution to the distribution of each layer in the DNN was calculated. As the measure of similarity we employed Kolmogorov Smirnov statistic subtracted from 1 since it is nonparametric and does not depend on the bin widths of the histogram very much. For each of the BMF and UCF for each of the rate and temporal coding, the similarities in the same regions in a single paper were averaged, and then the similarities in the same region in different papers were averaged (Extended Data Fig. 4). Averaging the 4 pairwise similarities (tBMF, tUCF, rBMF, and rUCF) derived the final layer-region pairwise similarity matrix. Since no distribution of tBMF was reported in AN, no distribution of rBMF was reported in AN, CN, or SOC, and no distribution of rUCF was reported in AN or CN, the similarities to them were set to 1 if there was no unit with definable BMF or UCF and set to 0 if otherwise. Also, for the regions other than those, the similarity was set to 0 if there was no unit with definable BMF or UCF.

### Evaluation of a pairwise similarity matrix

From a matrix of pairwise similarity, similarity of the entire cascade and that of each layer were calculated. We would like to evaluate the pairwise similarity matrix in a way that a DNN with its lower layers similar to the peripheral regions, its middle layers to the middle regions, and its higher layers to the central regions gets high score. To realize this concept of evaluation, we defined the similarity of the entire cascade, which we call cascade similarity, as the weighted mean of the pairwise similarity matrix (Extended Data Fig. 6). The weight at the position (<u>*i*</u>, *j*) was proportional to

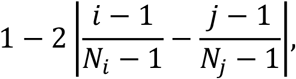

where *N_i_* and *N_j_* are the number of brain regions (= 7) and the number of the DNN layers, respectively. The weight was scaled so that the squared mean of the weight matrix was 1. The weight was maximum on the diagonal line and minimum on the top left and bottom right corners. Similarity of each layer, which we call layer-wise similarity, was defined as the mean taken in each layer.

### Control experiments

In the first control experiment, weights and biases were shuffled across units within each layer. The weights and biases were shuffled independently. In the second control experiment, category labels of the sounds in the training set were randomly shuffled. Validation set was not modified. The parameter update was conducted for the same number of iteration as the original non-random condition. In the third control experiment, the order of waveform samples in each sound was randomly shuffled, resulting in noise-like input waveform maintaining only the marginal distribution of the amplitudes. The fourth control experiment, the waveform following task, was to copy the amplitude value of the last timeframe of the input sound segment. To make the result directly comparable with those of the classification tasks, the target amplitude was quantized and the cost function was softmax cross entropy^62^. The waveform was nonlinearly transformed with a *μ*-law companding transformation before quantization^62^. The number of bins was equals to the number of sound categories in the original classification task.

## Acknowledgements

This work was supported by JSPS KAKENHI Grant Number JP15H05915 (Grant-in-Aid for Scientific Research on Innovative Areas “Innovative SHITSUKSAN Science and Technology”).

## Extended Data

**Extended Data Fig. 1.**
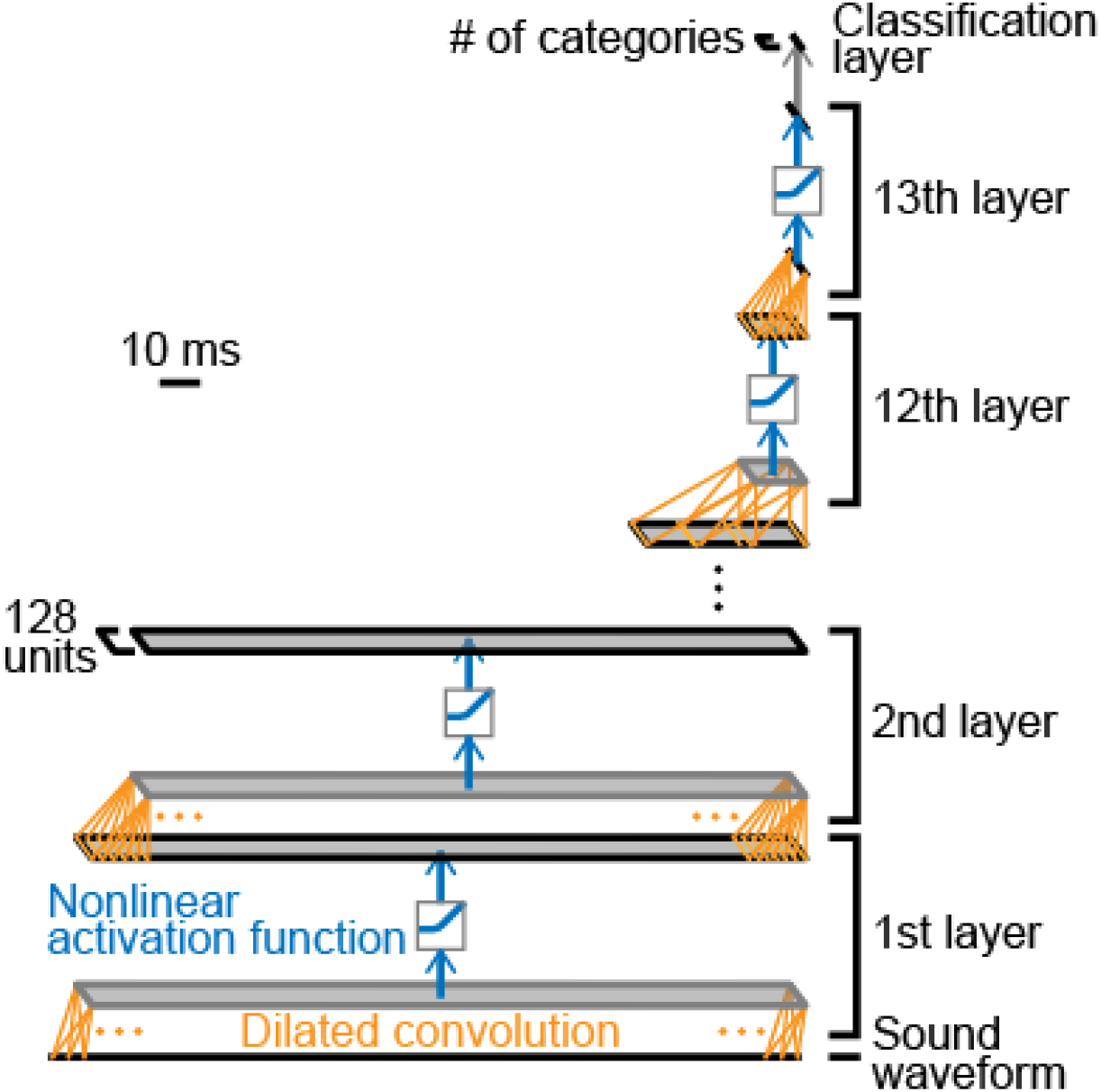
Architecture of the DNN. Our DNN consists of a stack of 1-dimentional dilated convolutional layers. The figure shows the architecture of the DNN for natural sounds. Each layer contains 128 units, and performs dilated convolution followed by nonlinear activation function. The 1st layer takes a raw sound waveform as an input, and the highest layer is connected to the classification layer, which was excluded from the analysis. The output category is the category assigned to the unit with maximum value. We tested multiple architectures with random filter and dilation length in each convolutional layer and selected the DNN which achieved the best classification accuracy on the novel dataset. The filter length and dilation length in all layers are shown in Extended Data Table 1. The number of layers and units in each layer was chosen in the pilot experiment. The activation function was the exponential linear unit.

**Extended Data Fig. 2.**
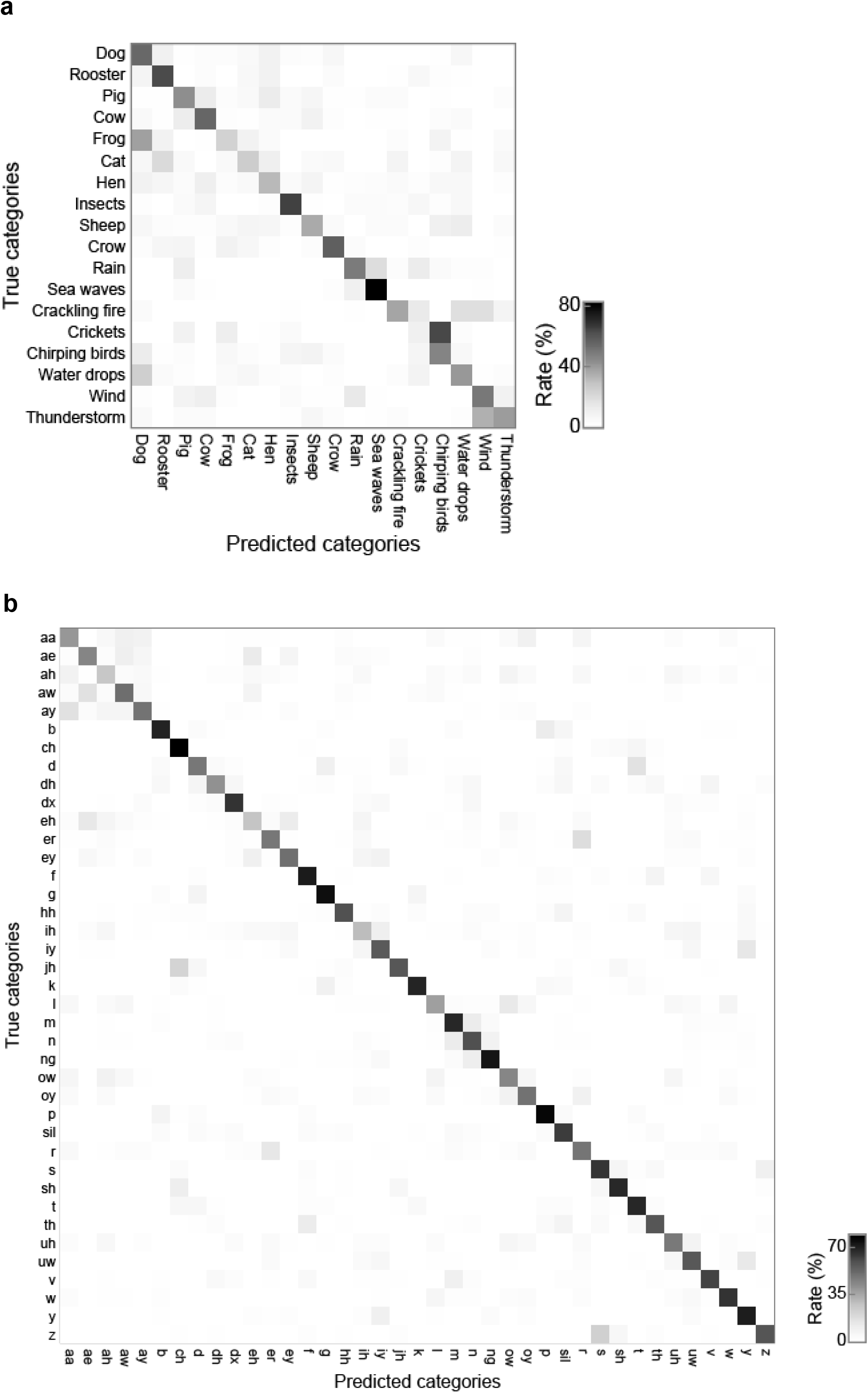
Confusion matrices of classification of the validation data. (a) Confusion matrix on the validation data of the non-human natural sounds. The number of categories are 18. (b) Confusion matrix on the validation data of the speech sounds. The number of categories are 39. Labels of true categories are shown in the ordinates and those of predicted categories are shown in the abscissas. The value in each cell is calculated as the fraction of timeframes classified to the particular category among the total timeframes with the true category. Cells with high classification rate are in the diagonal of the matrices, indicating the high classification accuracy. The classification accuracy was defined as the mean values in the diagonal of the matrix.

**Extended Data Fig. 3.**
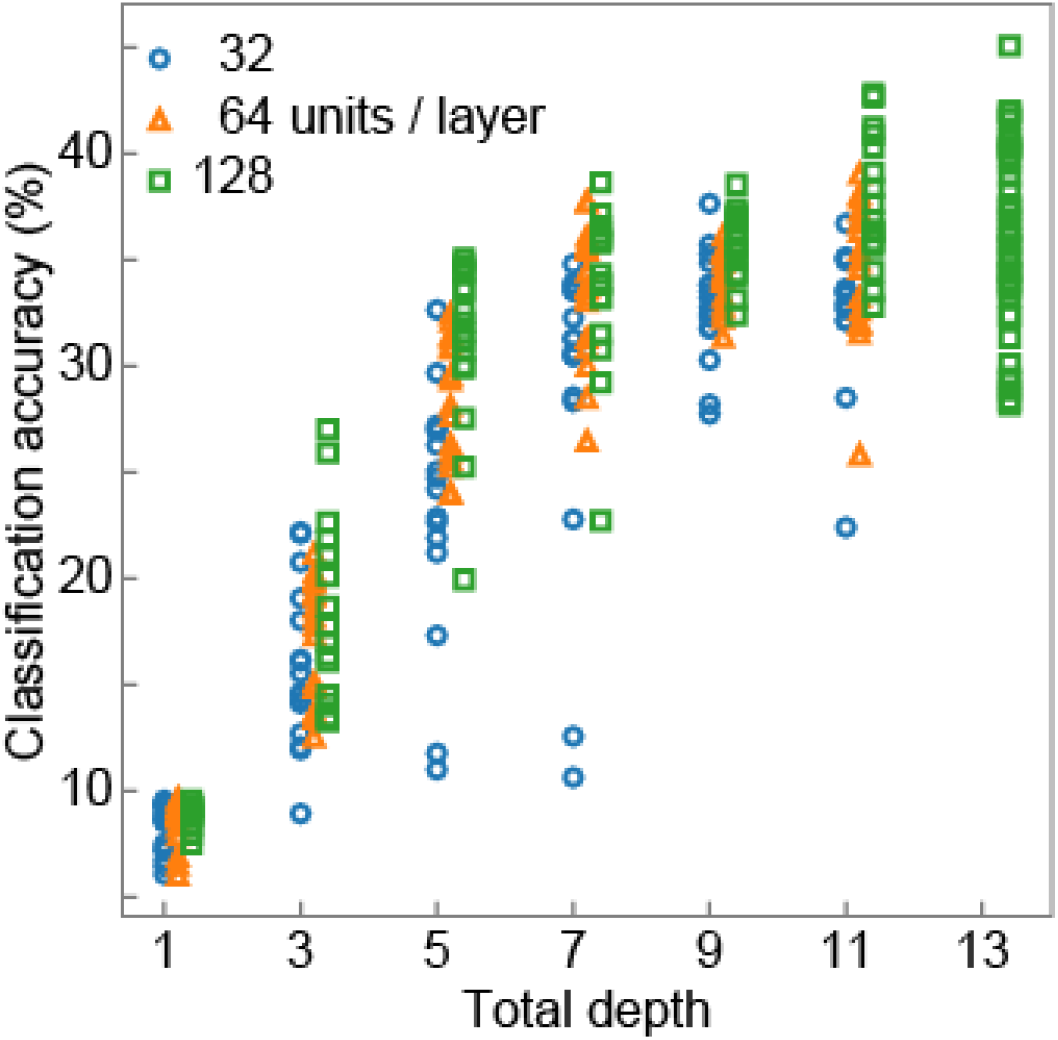
Importance of the deep cascade. Classification accuracy of DNNs with various number of layers with random filter and dilation length. DNNs with 1, 3, 5, 7, 9, 11, and 13 layers were tested. The number of tested channels were 32 (blue circles), 64 (orange triangles), and 128 (green squares). DNNs with 13 layers and 32 or 64 channels were not tested because they were excluded in the pilot study. The deeper the DNN, the higher the classification accuracy, seemingly saturating around the depth of 7. The result indicates the importance of the deep cascade at least as deep as 7 layers.

**Extended Data Fig. 4.**
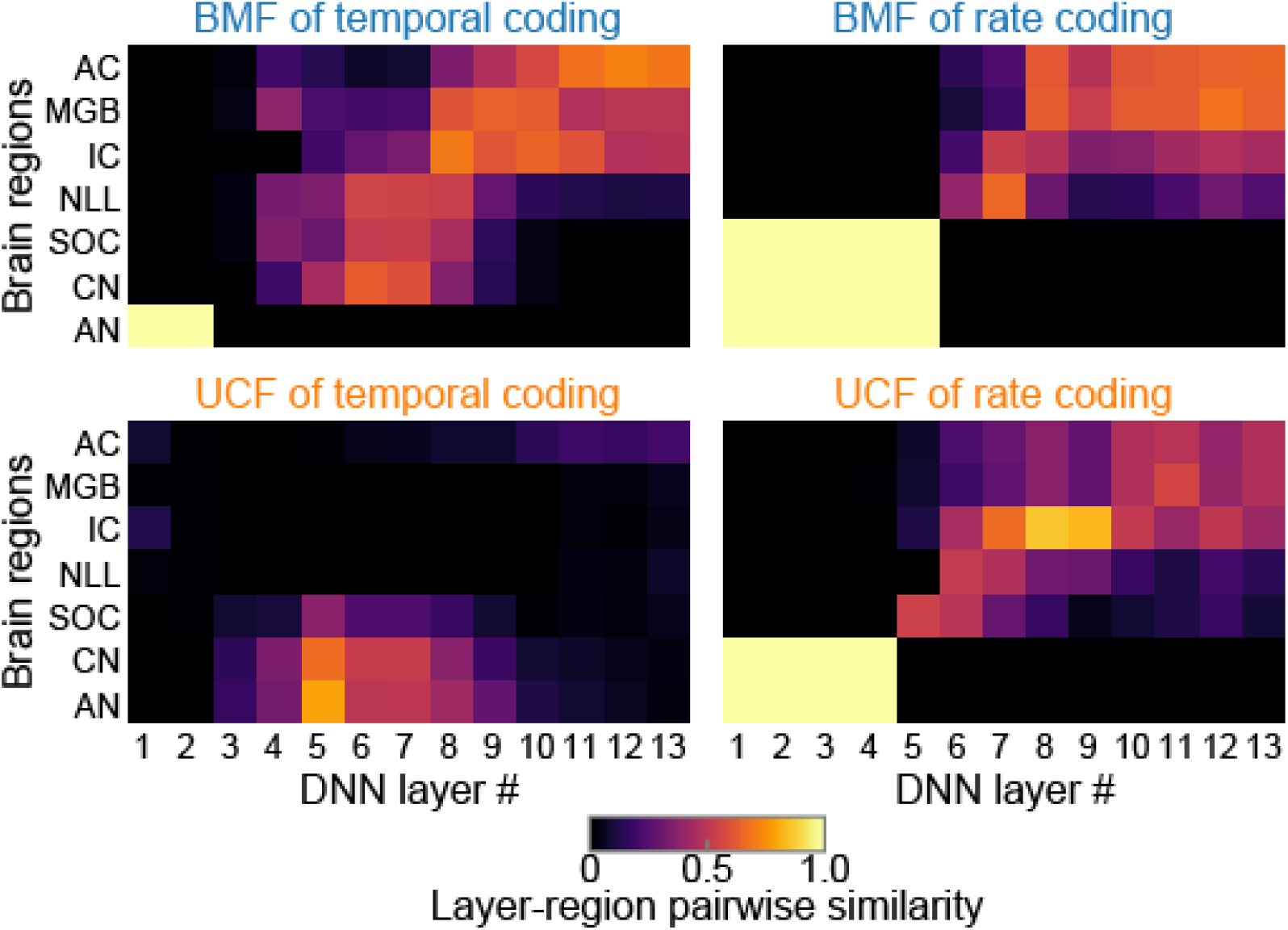
Layer-region pairwise similarity of each of the BMF and UCF of temporal and rate coding. Layer-region pairwise similarity of BMF (top panels) and UCF (bottom panels) of temporal (left panels) and rate (right panels) coding. The four pairwise similarities were averaged to yield the final layer-region pairwise similarity (Fig. 4d). In all of them, lower layers appeared to be similar to the peripheral regions and the higher layers to the central regions, although the similarities are not as smooth or clear as the averaged one.

**Extended Data Fig. 5.**
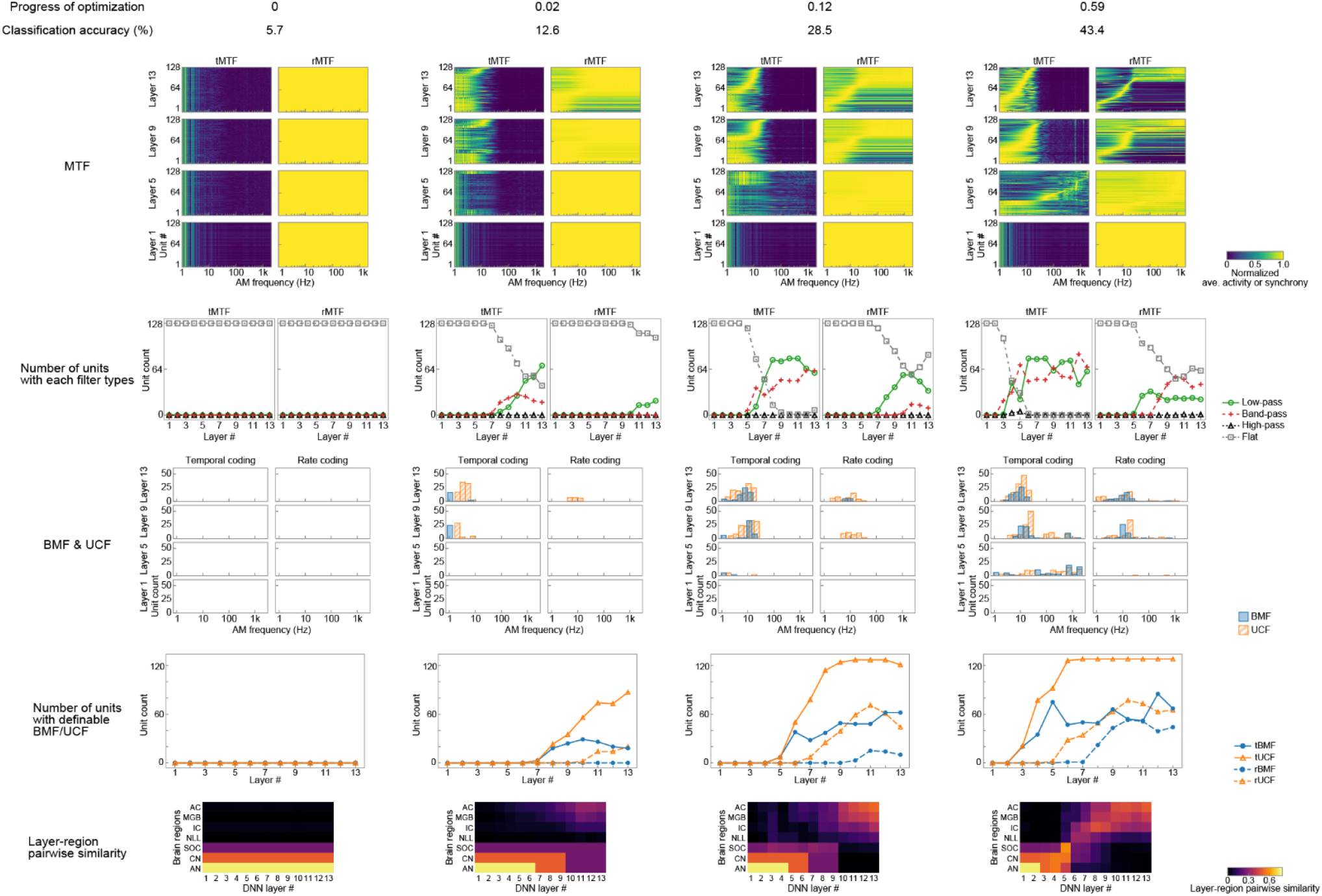
Development of AM representation in the DNN during optimization. From top to bottom: heatmaps of all tMTFs (left) and rMTFs (right) in layer 1, 5, 9, and 13 (as in Fig. 3c); the number of units with low-pass, band-pass, high-pass, and flat MTFs (as in Fig. 3b); histograms of BMFs and UCFs of temporal (left) and rate (right) coding (as in Fig. 4a); the number of units with definable tBMF, tUCF, rBMF, and rUCF (as in Fig. 4b); and layer-region pairwise similarity (as in Fig. 4d). The progress of the optimization and the classification accuracy is shown in the top of each column. Auditory-system-like AM tuning gradually emerged as optimization progressed.

**Extended Data Fig. 6.**
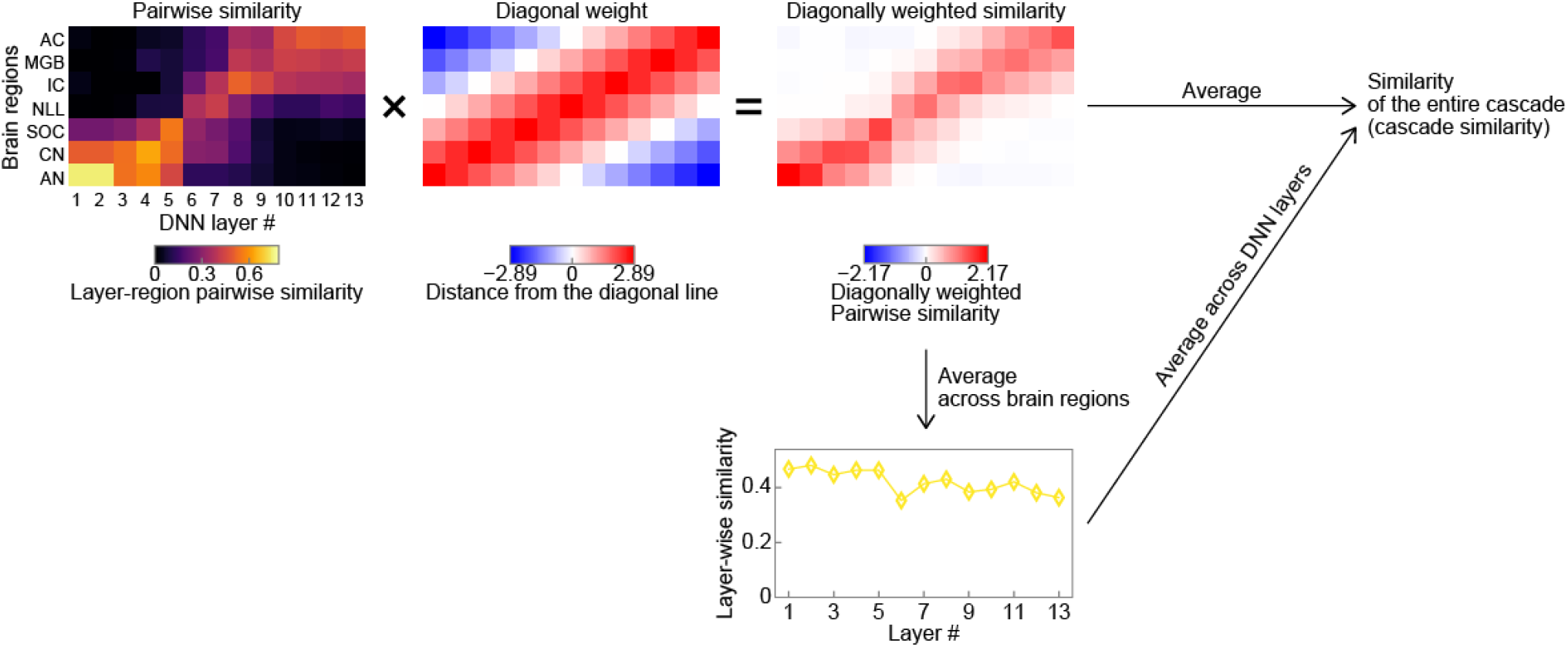
Calculation of the similarity of the entire cascade. Similarity of the entire cascade, which we call cascade similarity, was defined as the weighted mean of the pairwise similarity matrix. The weight was designed to be larger near the diagonal line and smaller in the left top and right bottom corners. The layer-wise similarity was defined as the mean calculated across brain regions within each layer.

**Extended Data Fig. 7.**
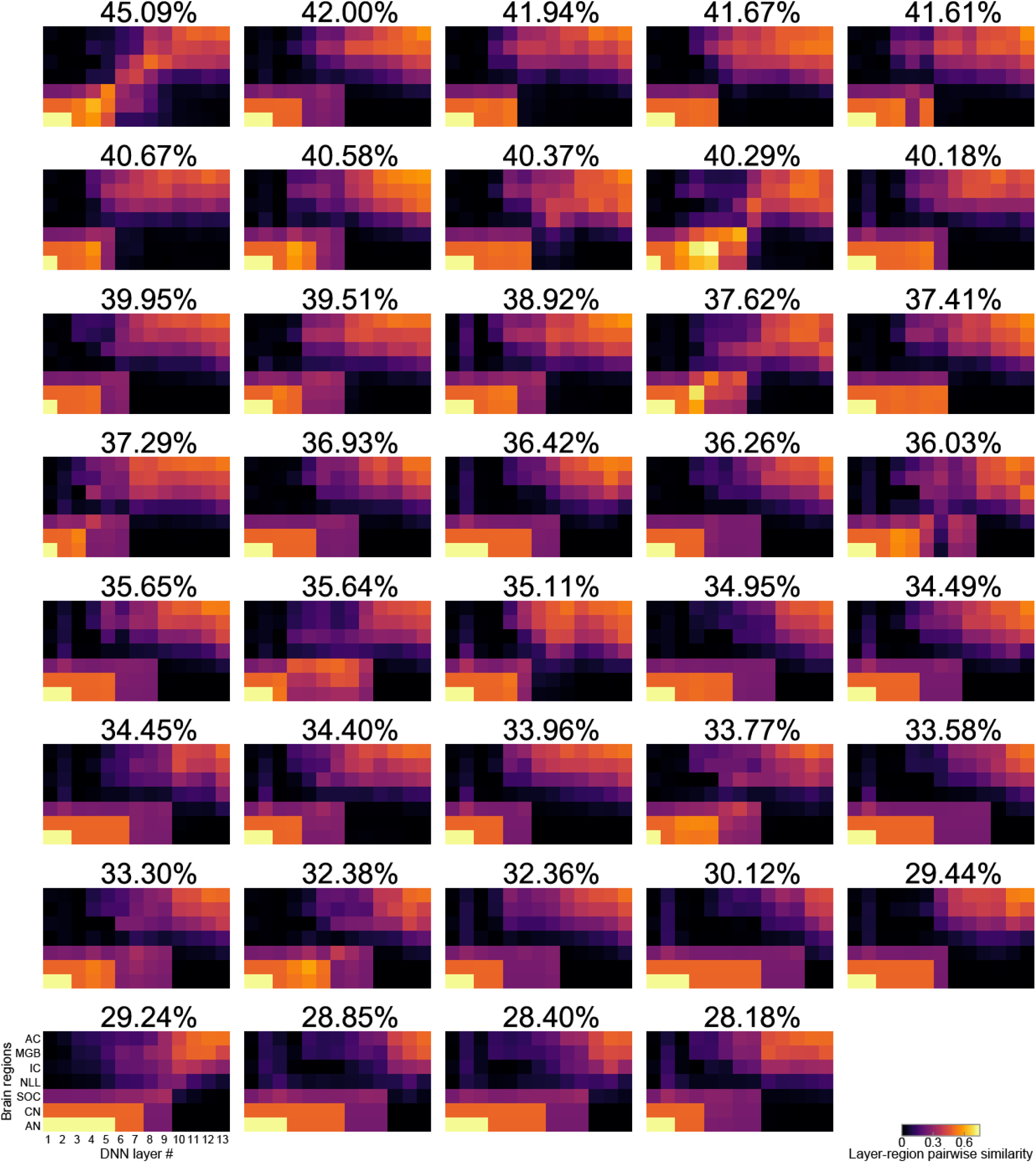
Layer-region pairwise similarity of the DNNs with various architectures. Heatmaps showing the layer-region pairwise similarity. The panels are sorted by the classification accuracy, shown in the top of each panel. The left top panel is identical to the one of Fig. 4d. Pairwise similarities in diagonal appeared larger in the DNNs with large classification performance.

**Extended Data Fig. 8.**
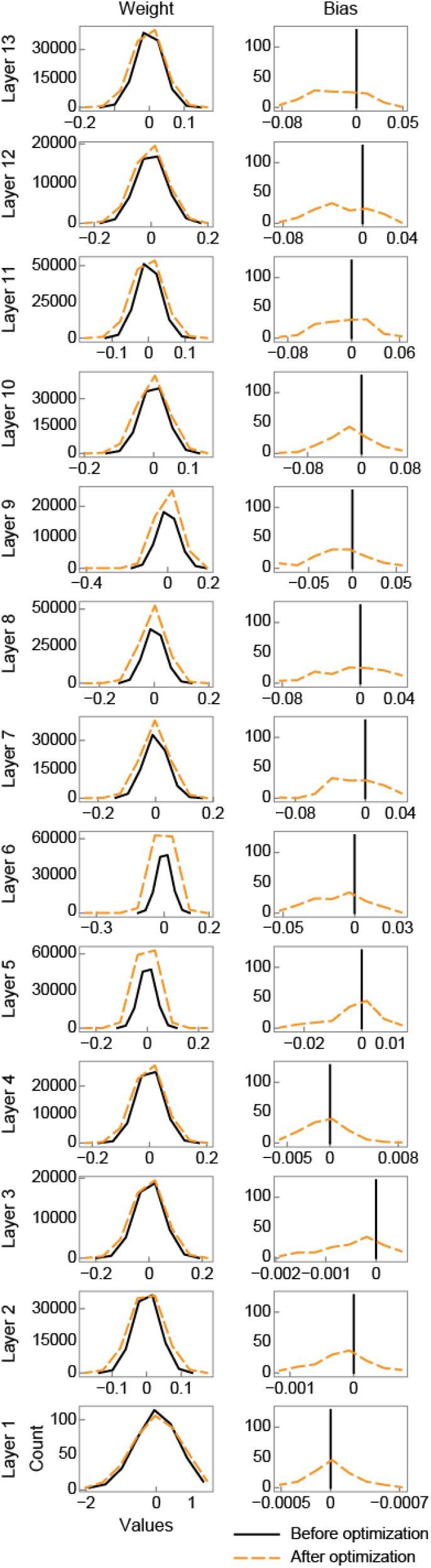
Distributions of the filter weights and biases before and after the optimization. Distributions of the filter weights (left panels) and biases (right panels) in each layer before (solid black lines) and after (dashed orange lines) the optimization. The layers are sorted vertically from bottom to top. In most layers the distribution of the filter weights appeared similar before and after the optimization. The distribution of the biases were totally different before and after since the biases before optimization are initialized to 0.

**Extended Data Fig. 9.**
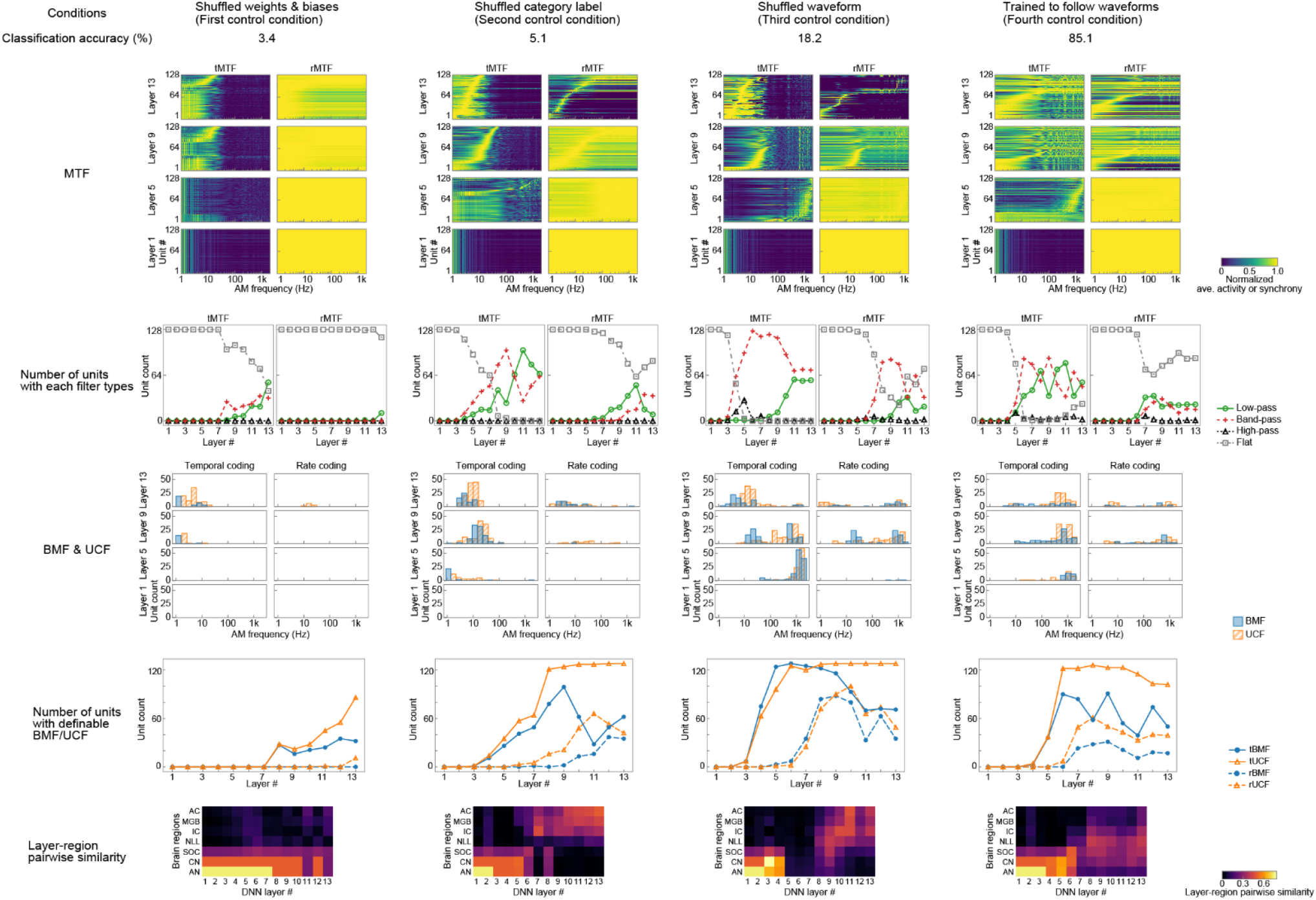
The lower layers were similar to the peripheral regions in all conditions. The middle layers were similar to the middle regions only in the fourth condition. The higher layers were similar to the central regions only in the second and third conditions. The results indicate different factors effecting AM representation in the different regions.

**Extended Data Fig. 10.**
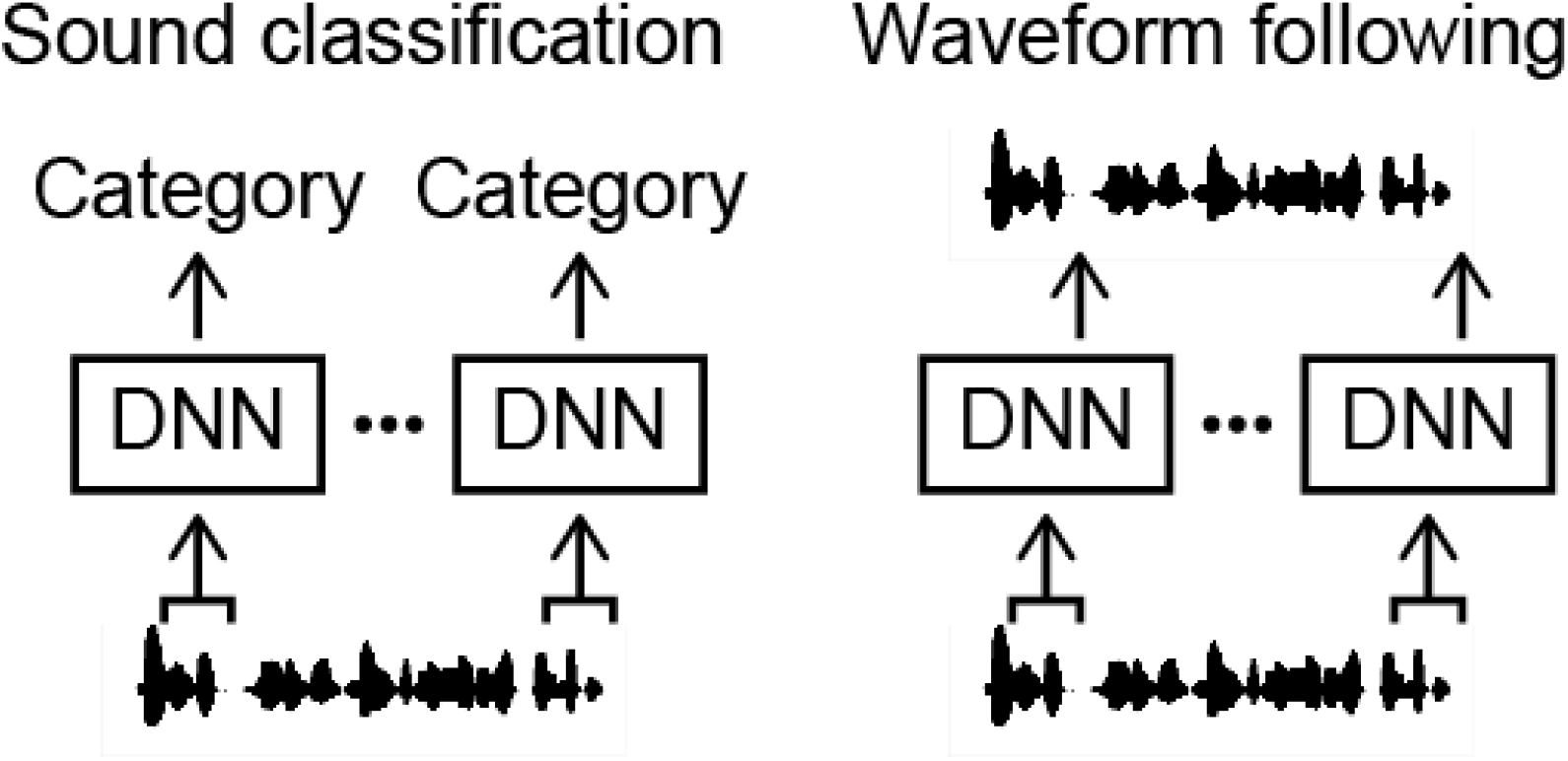
Schematic illustration of the classification task and the waveform following task. In the both tasks the DNN operated on a short sound segment. The sound classification task was to estimate the category of the input sound. The waveform following task was to copy the amplitude value of the last timeframe of the input segment.

**Extended Data Fig. 11.**
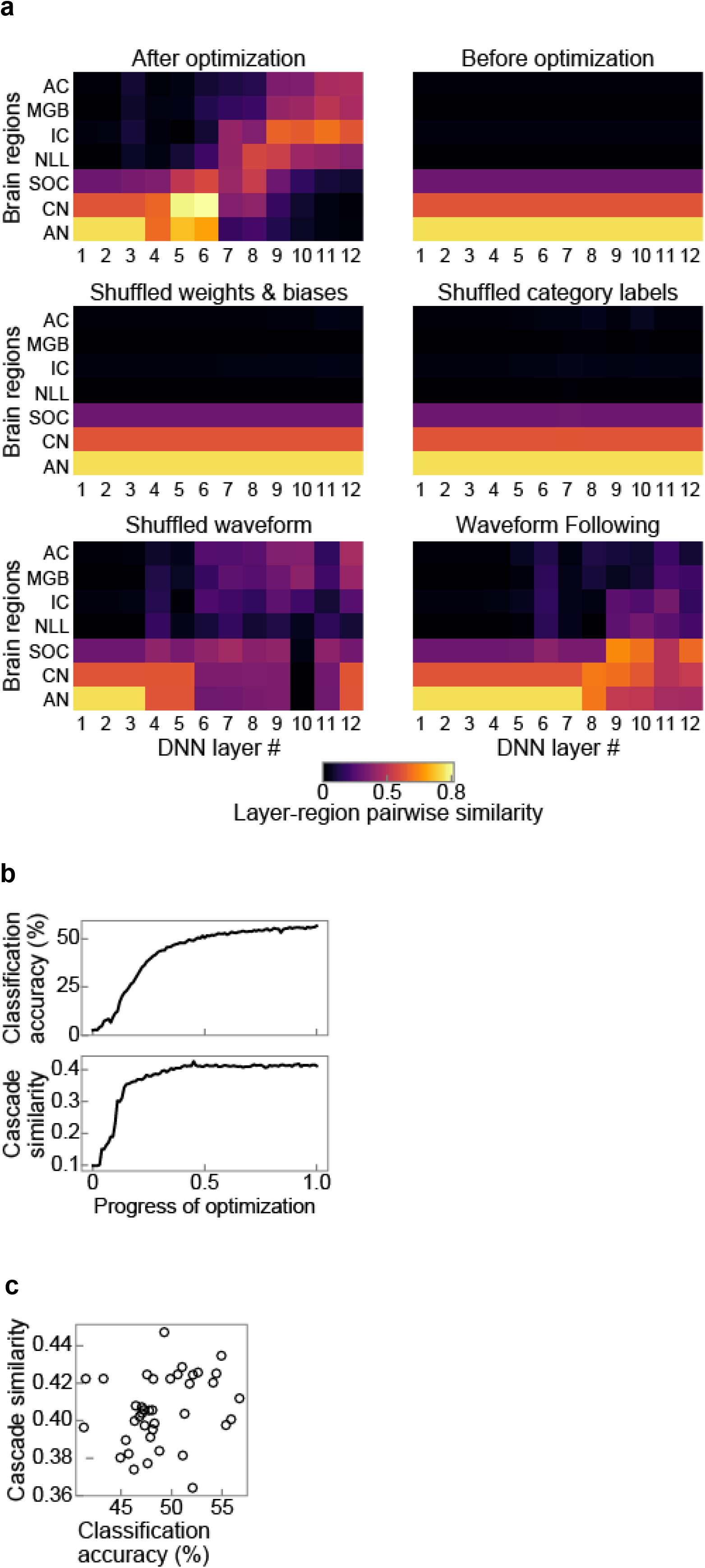
Similarity consistently emerges from the speech dataset. (a) Layer-region pairwise similarity after and before optimization, with shuffled weights and biases, trained on shuffled category labels and shuffled waveform, and of the waveform following task. Only did the DNN optimized for the classification task with natural data exhibited auditory-system-like AM representation. (b) The classification accuracy (top) and the cascade similarity (bottom) as functions of the progress of optimization. (c) The cascade similarities of the DNNs with various architectures, plotted against their classification accuracies. All results were consistent with the results obtained from the non-human natural sound.

**Extended Data Fig. 12.**
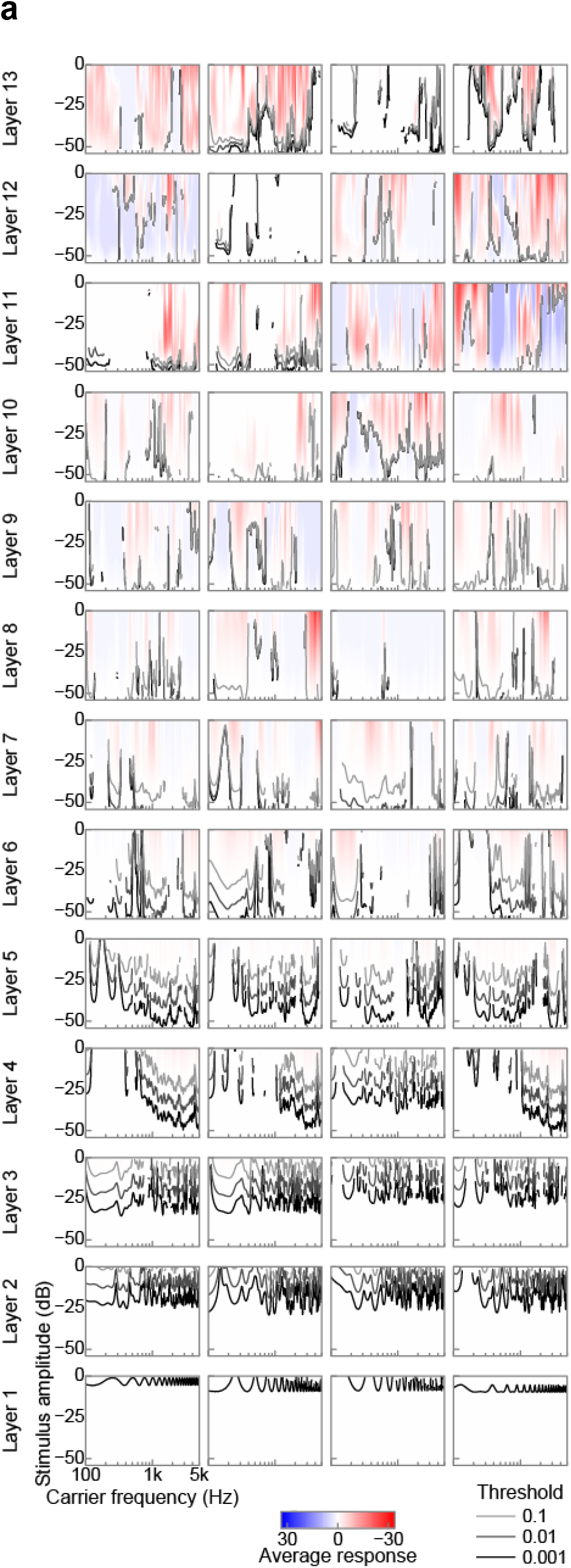
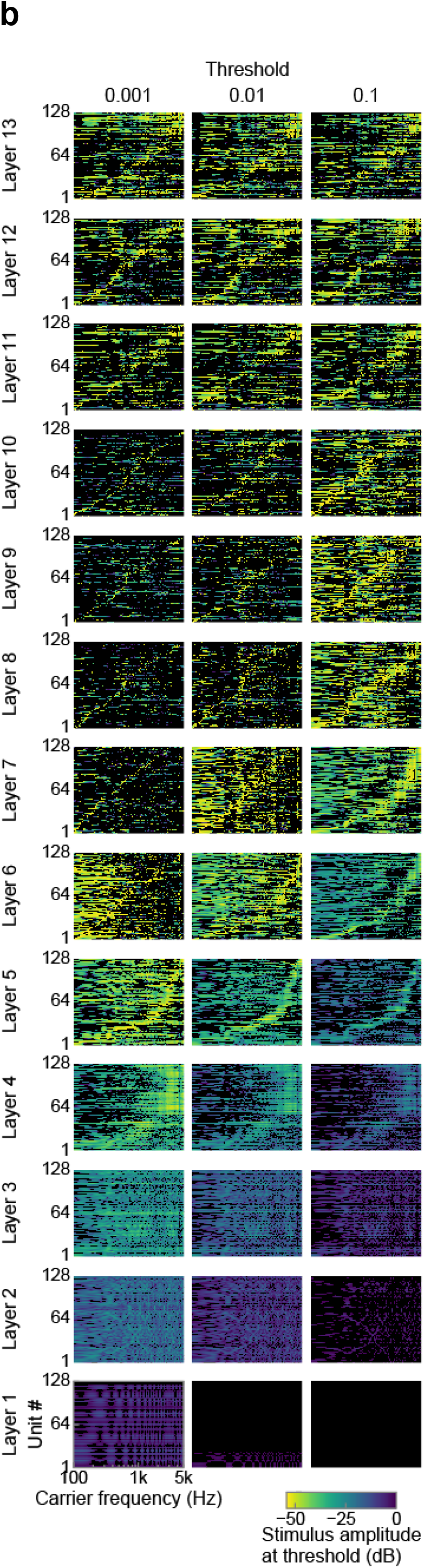
Tuning to carrier frequency. (a) Tuning to carrier frequency in 4 example units in each layer. Red and blue colour indicate larger and smaller response compared to the silent stimulus, respectively. White colour indicates the response equal to the silence. Black and grey lines show the frequency tuning curves, the minimum amplitude of the stimulus which induces larger response than the thresholds. The thresholds were 0.1 (light grey lines), 0.01 (dark grey lines), and 0.001 (black lines) above the response to the silence. Frequency tuning in the lower layers appeared monotonic along the stimulus amplitude, but some units in the higher layers shows non-monotonic response along the stimulus amplitude. The frequency tuning curves did not show clear single peaks. (b) Frequency tuning curve in all units in each layer. The curve for thresholds of 0.001 (left panels), 0.01 (middle panels), and 0.1 (right panels) above the response to the silence are shown. The units in each layer are sorted by the peak frequency of the tuning curves. Peaks in the frequency tuning curves in the middle layers appeared to cover wide range of the carrier frequency, but not in the lower and higher layers.

**Extended Data Table 1.** Architecture of the DNN.

15 Architecture of the DNN.
| Layer # | # channels | Dilation width | Filter width |
| --- | --- | --- | --- |
| 13 | 128 | 546 | 6 |
| 12 | 128 | 1189 | 3 |
| 11 | 128 | 1170 | 8 |
| 10 | 128 | 901 | 6 |
| 9 | 128 | 1129 | 3 |
| 8 | 128 | 1021 | 6 |
| 7 | 128 | 281 | 5 |
| 6 | 128 | 477 | 8 |
| 5 | 128 | 29 | 8 |
| 4 | 128 | 19 | 4 |
| 3 | 128 | 453 | 3 |
| 2 | 128 | 616 | 6 |
| 1 | 128 | 349 | 3 |

